# Accurate and rapid determination of metabolic flux by deep learning of isotope patterns

**DOI:** 10.1101/2023.11.06.565907

**Authors:** Richard C. Law, Kartikeya Pande, Samantha O’Keeffe, Glenn Nurwono, Rachel Ki, Pin-Kuang Lai, Junyoung O. Park

## Abstract

All life forms operate metabolism in constant flux. Metabolic fluxes offer a direct readout of cellular state, detailing the rates and driving forces of metabolic pathways. However, indirect, iterative solvers for mapping isotope patterns from tracing experiments onto metabolic fluxes leave much of cellular state uncharted. Here, we streamline metabolic flux quantitation by innovating a machine learning framework, ML-Flux, that deciphers complex isotope labeling patterns. We train neural networks using isotope pattern-flux pairs across central carbon metabolism from 26 key ^13^C-glucose, ^2^H-glucose, and ^13^C-glutamine tracers. ML-Flux takes variable-size isotope labeling patterns as input, imputes missing isotope patterns, and outputs mass-balanced metabolic fluxes. Computation of fluxes using ML-Flux is more accurate and faster than that of leading metabolic flux analysis software employing a least-squares method. Our biochemical networks and machine learning models constitute a curated and growing online knowledgebase of metabolic flux and free energy to democratize quantitative metabolic profiling.

---

Organisms employ dynamic networks of biochemical reactions to support proliferation, differentiation, homeostasis, cellular housekeeping, and bioproduct synthesis. Metabolism is one such network of pathways that provides cellular energy currency and biochemical building blocks. Metabolic fluxes represent the rates at which an organism operates these pathways. The culmination of biochemical knowledgebase^1–4^, analytical chemistry^5,6^, and mathematical modeling^7–11^ has established metabolic fluxes as a fundamental descriptor of cellular state in health and biotechnology^12–14^.

While metabolic fluxes indicate the dynamic state of an organism, they defy direct measurement because rates are intangible quantities. This challenge has become increasingly surmountable thanks to the development of stable isotope tracing and metabolic flux analysis (MFA) techniques^15–18^. As carbon forms the backbone of metabolites and biomass, tracing the fate of ^13^C from a ^13^C-labeled substrate to downstream metabolites reveals metabolic pathway utilization^19,20^. The accuracy of MFA relies on the use of cleverly chosen tracers that create differential isotope enrichment unique to each pathway and the measurement of isotope labeling patterns of metabolites located at the convergence of different pathways^21,22^. Using nuclear magnetic resonance spectroscopy (NMR)^23^ or mass spectrometry (MS)^24,25^ measurement of metabolite isotope labeling patterns and mathematical optimization, MFA searches for fluxes that simulate isotope patterns most consistent with experimental measurements. Isotope tracing and MFA tools are becoming increasingly integral to quantitative metabolic profiling^26–30^.

Despite its utility, MFA remains an expert method due to the need for judicious isotope tracer selection, custom metabolic model building, atom mapping across metabolic networks, and mathematical optimization. Furthermore, present MFA software using least-squares methods becomes computationally expensive with an increasing scope of metabolic networks and an increasing number of measured metabolites, restricting it to using only a handful of metabolites for simulation and data fitting out of hundreds of measurable metabolites^31^. To overcome these shortfalls, researchers require a simple mathematical function that accepts isotope labeling patterns as input and computes metabolic fluxes as output. For this function to be widely applicable, it needs to take in a variable-size input of isotope labeling patterns measured by different researchers studying divergent biological systems using different analytical instruments.

Here, we developed ML-Flux, a machine learning-based flux quantitation framework that maps isotope labeling patterns onto metabolic fluxes accurately and efficiently from nearly all conceivable isotope tracing experiments. We trained artificial neural networks (ANN) with neurons whose input signals of isotope labeling patterns were transformed into output signals of metabolic fluxes via synapse-like connections^32–34^. We further trained partial convolutional neural networks (PCNN) using convolution filters and binary masks to learn and impute missing isotope patterns from experimental measurements (*cf.* inpainting of images/matrices with missing pixels/elements)^35^. Integrating the two pre-trained neural network models, ML-Flux curtailed the time-consuming processes of constructing metabolic models and iterative flux estimations, thus streamlining the determination of metabolic fluxes and driving forces.

In computing metabolic fluxes, ML-Flux was consistently faster and >90% of the time more accurate than leading MFA software. Multiple isotope tracing in central carbon metabolism also revealed the unique advantages of ML-Flux: *i*) imputation of the isotope patterns of unmeasured metabolites (e.g., due to low abundance or instability); *ii*) inference of isotope labeling patterns in alternative tracer experiments; and *iii*) determination of metabolic flux and Gibbs free energy of reaction (ΔG)^36,37^. We released ML-Flux as an online resource to democratize flux quantitation (metabolicflux.org). The increased accessibility and knowledge of metabolic flux and free energy will accelerate sustainable biotechnology and therapeutic development by elucidating the dynamic state of biological systems.

## Results

### Fluxes create characteristic isotope labeling patterns

Our ability to quantify metabolic fluxes by isotope tracing relies on deciphering the relationship between metabolic fluxes and corresponding isotope labeling patterns. Tracing atoms from an isotope tracer (e.g., [1,2-^13^C_2_]-glucose) through metabolic pathways (e.g., glycolysis) leads to unique isotope labeling patterns of downstream metabolites (**Fig. 1a**). Given relative fluxes between two convergent pathways (e.g., upper glycolysis and pentose phosphate pathway, PPP), one can compute the isotope pattern of a metabolite at the merge point (e.g., glyceraldehyde-3-phosphate, GAP) by linear combination (**Fig. 1a**). Therefore, simulating isotope labeling patterns from known metabolic fluxes uses straightforward linear algebra (**Fig. 1b**).

**Figure 1.**
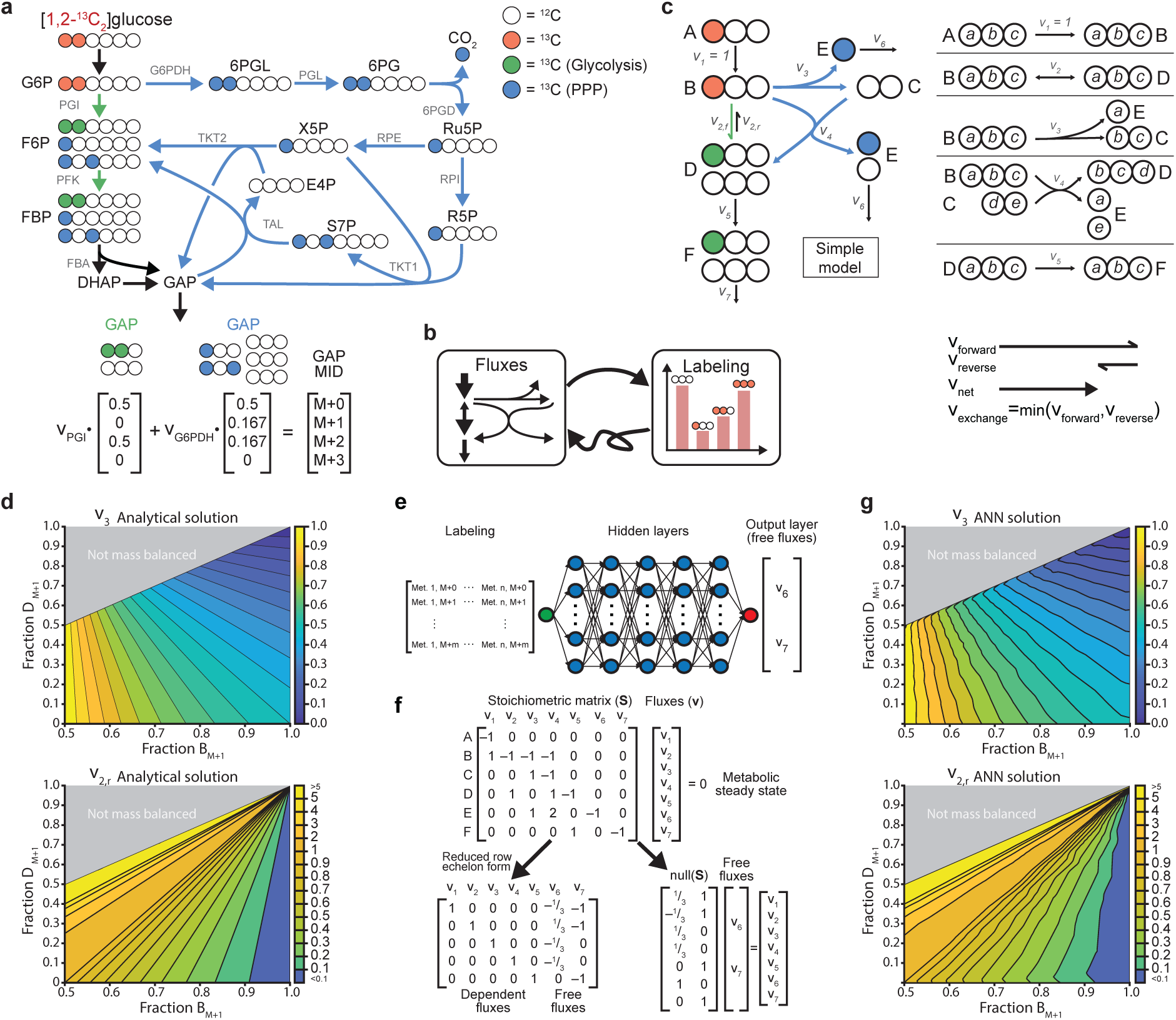
Isotope tracers imprint metabolic fluxes on metabolites in the form of isotope patterns. **a,** Tracing [1,2-^13^C_2_]-glucose discerns relative usage of glycolysis and the pentose phosphate pathway (PPP). The mass isotopomer distribution (MID) of glyceraldehyde-3-phosphate (GAP) is a linear combination of pathway-specific isotopologues weighted by fluxes. **b,** Simulating metabolite MIDs from an isotope tracer given metabolic fluxes is a straightforward process, but the inverse process of quantifying fluxes given MIDs is convoluted and indirect. **c,** A simplified model of glycolysis and PPP illustrates how atomic transitions impact isotope labeling patterns. Feeding [1-^13^C_1_]-A into the system results in unique isotope patterns as a function of v_3_ and reverse flux v_2,r_. **d,** Analytical solutions for v_3_ and v_2,r_ were solved as functions of B and D isotope labeling. Each axis represents the fraction of the singly labeled (M+1) isotopomer, while the color scale represents the flux value. **e,** An artificial neural network (ANN) was trained to take isotope labeling patterns as input and predict free fluxes. **f,** The remaining dependent fluxes were calculated as a linear combination of the free fluxes. **g,** The trained ANN model reproduced the isotope pattern-to-flux relationship identified in the analytical solutions from panel **d**.

The inverse process of mapping isotope labeling patterns to fluxes is nonlinear, convoluted, and often unknown (**Fig. 1b**). As a result, fluxes are conventionally determined by recursive simulation. Only in the simplest of cases can fluxes be calculated directly using an analytical relationship between isotope labeling patterns and fluxes. We demonstrated this in a simple toy metabolic model mimicking upper glycolysis and the PPP^15^ that contains two free net fluxes (v_1_ and v_3_) and one exchange flux (v_2,r_) (**Fig. 1c**). We obtained an analytical solution to the fluxes by tracing molecule ‘A’ harboring a heavy isotope in its first position (i.e., [1-^13^C_1_]) and imposing steady-state mass balance on the ensuing isotopologues:

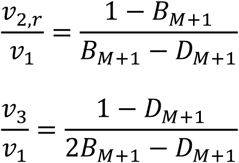

B_M+1_ and D_M+1_ are the fractions of the singly labeled B and D (i.e., M+1 isotopomers). These solutions showed that fluxes are (non-linear) functions of isotope labeling patterns and that isotope patterns are characteristic of underlying fluxes (**Fig. 1d**).

### Neural networks predict fluxes from teachable isotope patterns

To test if flux-dependent isotope patterns can be taught, we trained an ANN with isotope labeling patterns simulated using the simple toy metabolic model and the [1-^13^C_1_]-A tracer. Principal component analysis of isotope patterns revealed unique features that we hypothesized deep learning could use to formulate accurate isotope pattern-to-flux relationships (**Extended Data Fig. 1a,b**). ANNs were trained using fluxes sampled from multiple distributions to determine the optimal dataset for learning (**Extended Data Fig. 1c,d**). Log-uniform flux sampling resulted in the best ANN model. It predicted the free fluxes accurately throughout all ranges of mass isotopomer distribution (MID) input (**Fig. 1e**). The remaining fluxes were obtained by multiplying the free fluxes by the basis of the null space of the metabolic model (**Fig. 1f**). The resulting ANN model orthogonally mapped isotope patterns to fluxes nearly identically to the analytical solution (**Fig. 1g**).

Solving isotope pattern-to-flux functions becomes increasingly challenging as the size of the metabolic network (i.e., number of reactions and atoms) and number of isotope tracers used grows. We set out to test if isotope pattern-to-flux relationships could still be taught for more complex metabolic networks without an apparent analytical solution for linking isotope labeling patterns to fluxes. We developed models of upper and full glycolysis, glycolysis and the PPP (GlyPPP), and central carbon metabolism (CCM) (**Fig. 2a**). To obtain training data, we simulated isotope patterns using 24 combinations of commercially available ^13^C-glucose, [5-^2^H_1_]-glucose, and ^13^C-glutamine (**Supplementary Table 1**) across a physiological flux space (**Supplementary Tables 2-5**). For the simple toy model, we simulated isotope patterns from all six nontrivial isotope tracers (i.e., all but the uniformly labeled and the uniformly unlabeled ones). For each of the five metabolic models, we trained an ANN to compute metabolic fluxes from the MIDs of all constituent metabolites.

**Figure 2.**
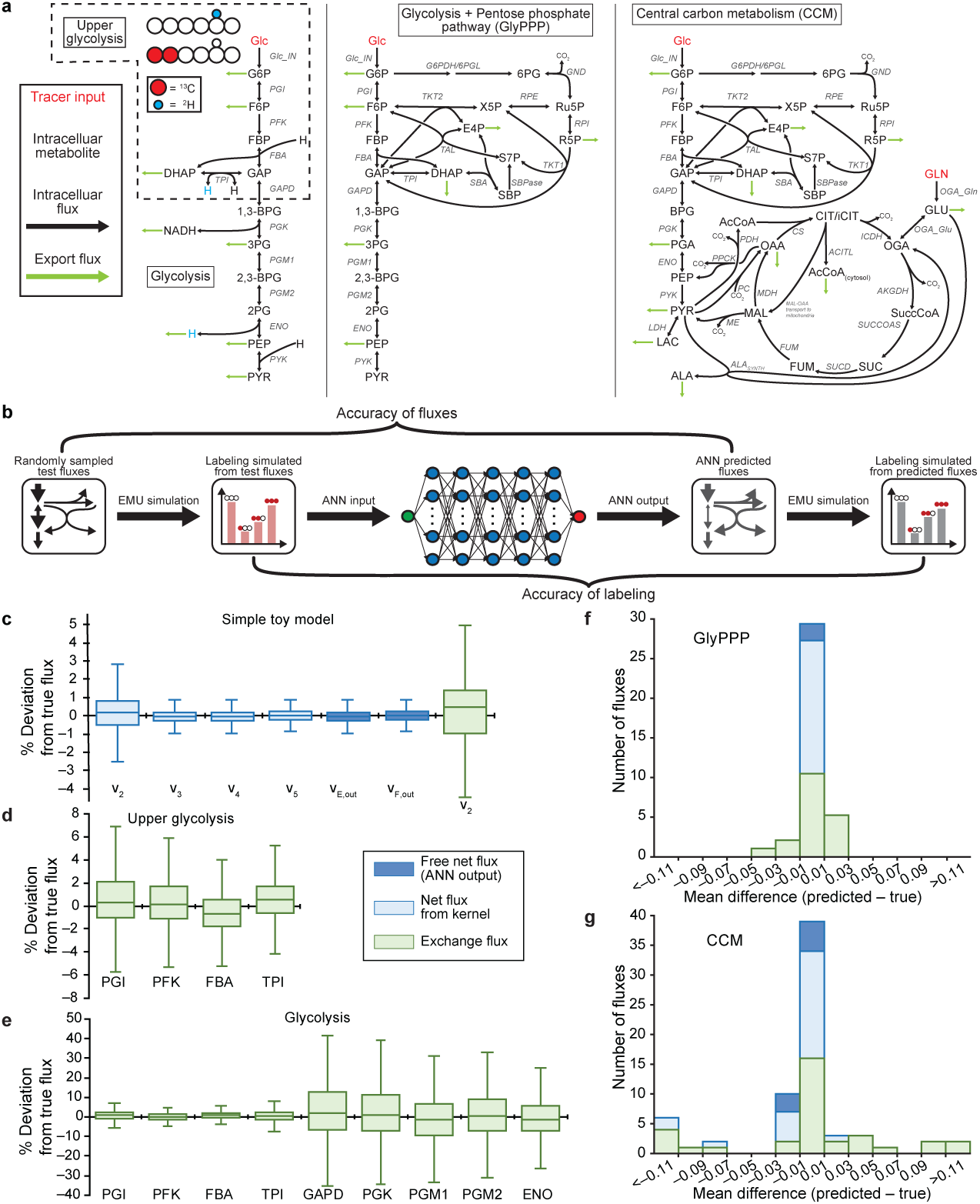
Artificial neural networks decipher the relationship between metabolite isotope labeling patterns and reaction fluxes. **a,** ANNs were trained to predict fluxes in four metabolic networks. Each metabolic model was used to simulate isotope labeling patterns from randomly sampled fluxes using multiple isotope tracers. **b,** Isotope labeling data from test fluxes were inputted into ANNs to predict fluxes. Predicted fluxes were then used to simulate isotope labeling patterns. Comparing test and predicted fluxes described ANN flux prediction accuracy. The accuracy of labeling was also evaluated by comparing isotope labeling patterns simulated from test and predicted fluxes. **c-e,** Free net (dark blue) and exchange (green) fluxes were predicted from testing data for each metabolic model. Dependent net fluxes (light blue) were computed from free net fluxes. The distribution of errors associated with flux predictions were plotted for (**c**) the simple toy, (**d**) upper glycolysis, and (**e**) glycolysis models. Each box shows the three quartiles, and whiskers extend to the minimum and maximum values within 1.5-fold of the interquartile range (n=10,000). **f-g,** The mean error of each reaction flux across all testing data was computed for the reactions in (**f**) GlyPPP and (**g**) CCM models (n=100,000 and 11,708, respectively).

We assessed the performance of the trained ANN models using reserved testing data (**Fig. 2b**). A higher flux prediction accuracy generally corresponded to a greater agreement between test isotope patterns and simulated isotope patterns from the predicted fluxes (**Extended Data Fig. 2**). In the simple toy model, both net and exchange fluxes were predicted within 5% error across all testing data (**Fig. 2c**). In the glycolysis models, which were trained with [1,2-^13^C_2_]-glucose and [5-^2^H_1_]-glucose tracers, flux predictions were within 10% of test data for the exchange fluxes in upper glycolytic reactions (**Fig. 2d**). Exchange fluxes in lower glycolytic reactions displayed prediction errors with a ±∼10% interquartile range (**Fig. 2e**). A high reversibility of the triose phosphate isomerase (TPI) reaction (i.e., a high exchange flux) lowered the sensitivity of exchange fluxes in lower glycolysis to deuterium labeling fractions from [5-^2^H_1_]-glucose (**Extended Data Fig. 3a**). With a lower TPI reversibility (i.e., exchange flux≤2), the predictions of lower glycolytic exchange fluxes became more accurate with an interquartile range of ±∼5% (**Extended Data Fig. 3b**). For the GlyPPP model, all flux prediction errors fell within ±0.03 flux units (fluxes normalized to glucose uptake) (**Fig. 2f**). For the CCM model, 85% of CCM flux predictions were accurate within ±0.05 flux units (**Fig. 2g**). Flux predictions were robust to small variances in isotope labeling measurement that may arise from instrument measurement error (**Extended Data Fig. 3c-f**). Regression analysis showed that the coefficient of determination (R^2^) for flux predictions ranged from 0.9 to 1 with the slopes between prediction and truth having a strong central tendency around 1 (**Extended Data Fig. 4a**). Reduced chi-squared (χ^2^) test showed that 95% of flux predictions were statistically acceptable with a relative standard deviation of 0.10 for net fluxes and 0.68 for exchange fluxes (**Extended Data Fig. 4b**). Based on the distributions of prediction errors in the test data, we derived the standard errors for individual flux predictions (**Supplementary Table 6**). The goodness-of-fit analyses validated the ability of our ANNs to accurately compute nearly all net fluxes and many exchange fluxes.

### Inpainting permits variable-size input of isotope patterns for flux determination

An obstacle to the widespread adoption of ANN models for flux analysis was their uniform input requirement. The ANN architecture requires a rigid input structure consistent with its training data (i.e., the isotope labeling patterns of the full set of metabolites). However, metabolite measurements are seldom complete due to varying analyte abundance, stability, environment, and instrumentation. Furthermore, experimentalists may employ various tracers that harbor isotopes at different positions^38–41^. We sought to bridge the gap between experimental measurement and ANN input by imputation. To bring partial isotope pattern measurements to complete input data suitable for ANN, we took two approaches: K-nearest neighbors (KNN) regression^42^ and inpainting based on partial convolutional neural networks (PCNN)^35^.

In KNN regression, the unknown (unmeasured) isotope labeling patterns were assigned the Euclidean distance-weighted mean of corresponding values from its three nearest neighbors (K=3) in the training data (**Fig. 3a**). To test the KNN approach, we generated partially masked isotope patterns from testing data to mimic typical experiments with incomplete metabolite measurements resulting from one to a few randomly selected isotope tracers at a time. The mean absolute errors of predicted isotope labeling fractions were less than 0.01 for glycolysis models for computing driving forces and less than 0.0005 for the models of central carbon metabolism (**Fig. 3b**).

**Figure 3.**
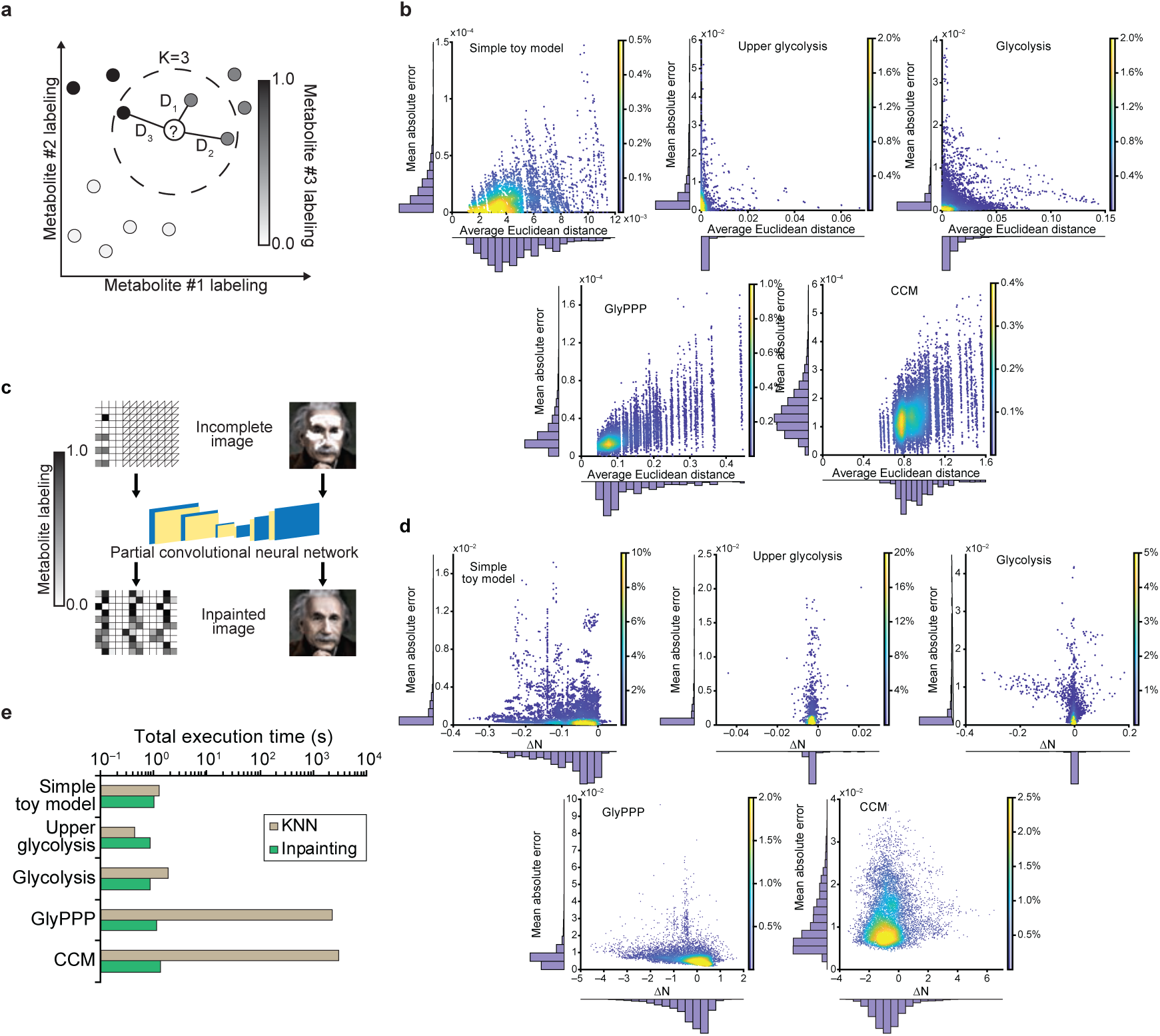
Imputation and inpainting algorithms complete variably missing isotope labeling pattern information. **a,** The known parts of an incomplete isotope labeling pattern set were compared with the corresponding parts of complete isotope labeling pattern sets to identify the three nearest neighbors by Euclidean distance. The missing parts of the incomplete isotope patterns were imputed as the weighted means of the corresponding known isotope patterns of three nearest neighbors. **b,** For each metabolic network, 100 randomly generated masks partially covered each of 100 complete isotope patterns to show only a subset of metabolites for 1-3 tracer experiments. These masked datasets were imputed with KNN regression. The mean absolute errors of predicted isotope patterns were shown along with the average Euclidean distances to three nearest neighbors. **c,** MIDs represented as a matrix resembled pixels in a black-and-white image. Missing elements in an MID matrix were predicted akin to how missing pixels in an image are restored by inpainting by a partial convolutional neural network (PCNN). **d,** For each metabolic network’s PCNN, the mean absolute errors of predicted isotope labeling patterns were shown along with the plausibility score (ΔN), the difference between the grand sum of the inpainted MID matrix and the total number of measured metabolite instances. **e,** The KNN regression and PCNN inpainting were benchmarked by their total execution time to impute missing information in one set of incomplete isotope labeling patterns.

In PCNN-based inpainting, the isotope pattern matrix was treated as a grayscale image with pixels corresponding to fractions of mass isotopomers between 0 and 1 (**Fig. 3c**). The PCNN inpainting model was trained to verisimilarly fill in the irregular missing regions of the isotope pattern matrices covered by random binary masks. The binary masks exposed the isotope patterns for a random subset of metabolites resulting from one to a few randomly selected isotope tracers at a time. To test the plausibility of inpainted isotope pattern matrices, we developed a simple metric ΔN, the difference between the grand sum of the inpainted MID matrix and the total number of measured metabolite instances, since the sum of the labeling fractions for each metabolite should be 1. As ΔN approaches 0, plausibility increases. The trained PCNN inpainting model resulted in predictions of missing data that were highly plausible and accurate with mean absolute errors less than 0.02 (**Fig. 3d**). In addition to accuracy, a major benefit of the PCNN model was its fast computation time that was invariant with respect to the complexity of the underlying metabolic networks (**Fig. 3e**).

### Two-stage machine learning streamlines metabolic flux determination

We integrated the PCNN and ANN models to map variable-size input of isotope pattern measurements onto metabolic fluxes (**Fig. 4a**). We quantified the accuracy of flux predictions by the two-stage ML using partially known isotope patterns resulting from a small subset of available isotope tracers. In the simple toy model and two glycolysis models, 40-95% of isotope patterns were masked. The median error of the PGI exchange flux rose from 1% to 10% in the glycolysis model, whereas the TPI exchange reaction was less affected by the inpainted isotope patterns, increasing from 3% to 8% (**Fig. 4b**). The flux prediction accuracy depended on whether isotope labeling data from key metabolites were input (**Extended Data Fig. 5**). In the GlyPPP and CCM models, two-stage ML predictions starting from >97%-masked isotope patterns data reduced the number of fluxes predicted with <10% error, and the errors for some exchange fluxes increased by up to 20% (**Fig. 4c**, **Extended Data Fig. 6**, and **Supplementary Table 7**). These observations suggested the tradeoff between the flexible isotope pattern input and the flux prediction accuracy.

**Figure 4.**
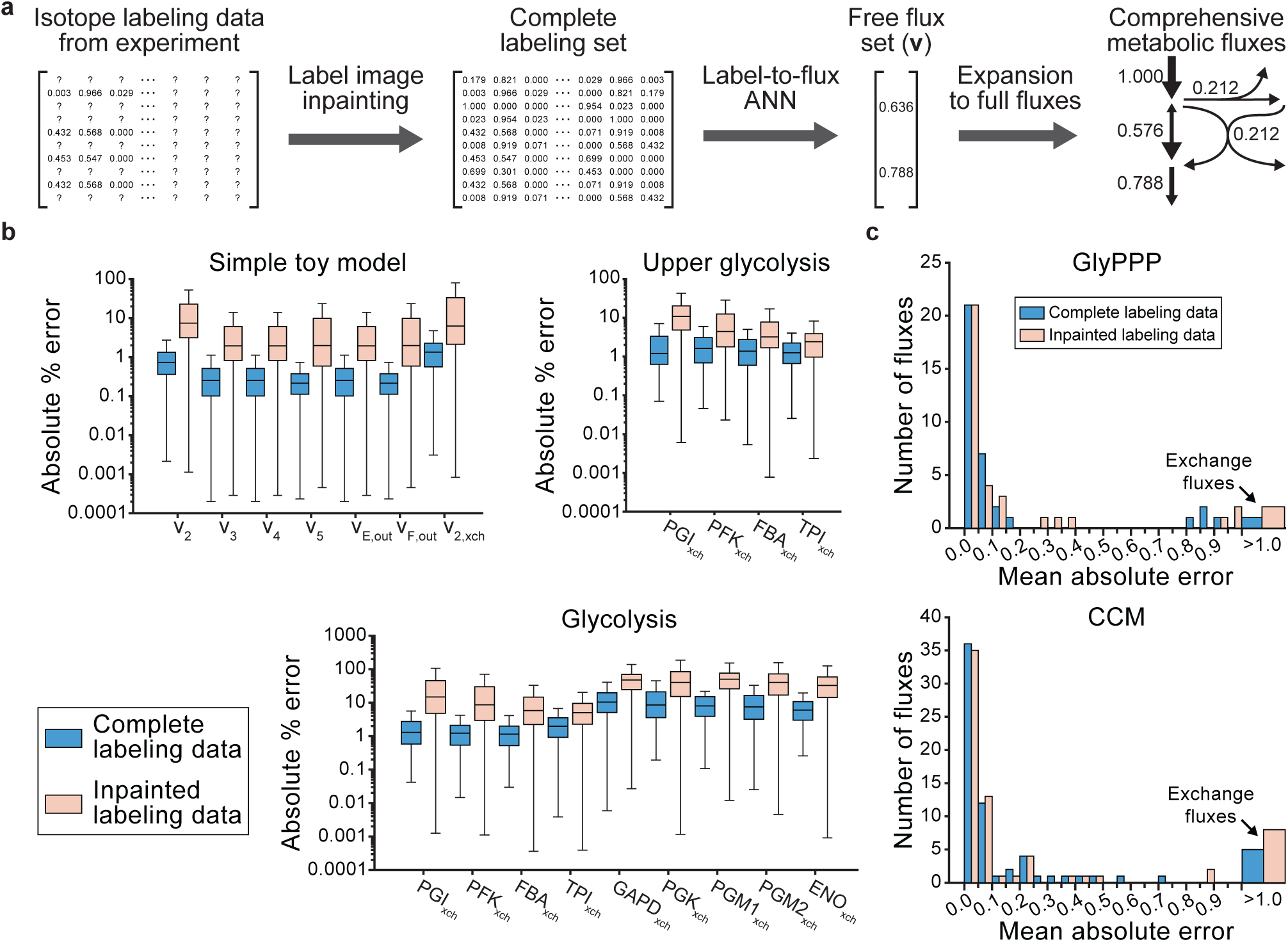
Integrated ML models produce accurate fluxes from variable-size input isotope patterns. **a,** PCNN inpainting and ANN flux prediction were integrated to convert variable-size metabolite isotope labeling patterns into free fluxes, which were expanded to a comprehensive flux map using linear algebra. **b,** Fluxes were predicted either from complete isotope pattern information or masked versions of the same datasets after the PCNN inpainting. The distribution of their absolute percent errors was plotted for the simple toy, upper glycolysis, and glycolysis models. Each box shows the three quartiles, and whiskers extend to the minimum and maximum values within 1.5-fold of the interquartile range (n=100 for complete isotope labeling patterns, n=10,000 for simple toy and glycolysis masked isotope labeling patterns, and n=3,100 for upper glycolysis masked isotope labeling patterns). **c,** The distributions of mean absolute errors of fluxes in the GlyPPP and CCM models were plotted for complete and inpainted isotope labeling patterns (n=100 for complete isotope labeling patterns, n=10,000 for masked isotope labeling patterns).

Select tracers are commonly used for their ability to accurately capture fluxes^43,44^. To identify which isotope tracers best potentiate ML-Flux to accurately predict fluxes, we subjected the two-stage ML model for central carbon metabolism to single tracing, dual tracing (i.e., one ^13^C-glucose and one ^13^C-glutamine tracers in a single experiment), and parallel tracing (i.e., two ^13^C glucose tracers in duplicate experiments). Dual tracing of [4-^13^C_1_]-glucose and [U-^13^C_5_]-glutamine displayed the highest net flux prediction accuracy throughout central carbon metabolism (**Extended Data Fig. 7a** and **Supplementary Table 8**), outperforming pervasive parallel tracing experiments using [1,2-^13^C_2_]-glucose and [U-^13^C_5_]-glutamine (**Extended Data Fig. 7b**). Therefore, with judicious tracer selection, two-stage ML models with minimal, variable-size isotope pattern input accurately predicted fluxes in accordance with goodness-of-fit analysis.

### Neural networks predict metabolic driving forces from isotope patterns

Accurate prediction of both net and exchange fluxes yielded additional insights into thermodynamic driving force, enzyme efficiency, and pathway flux control. According to a fundamental thermodynamic principle, Gibbs free energy of reaction (ΔG) is log-proportional to reaction reversibility (defined as the ratio of reverse to forward fluxes, *v_reverse_*/*v_forward_*)^4,36,37^:

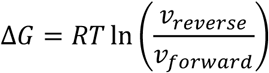

R is the universal gas constant, and T is temperature in Kelvin. Strongly forward-driven reactions with ΔG<<0 kJ/mol often correspond to rate-controlling pathway steps^45^, whereas near-equilibrium reactions with –2<ΔG<0 kJ/mol imply highly adaptivity and spare enzyme capacity^4^. Select isotope tracers reveal the extent of reaction reversibility and thus ΔG (e.g., [1,2-^13^C_2_]-glucose and [5-^2^H_1_]-glucose for glycolytic reactions) because the reverse reaction dilutes isotope enrichment in metabolites (**Fig. 5a**)^46^.

**Figure 5.**
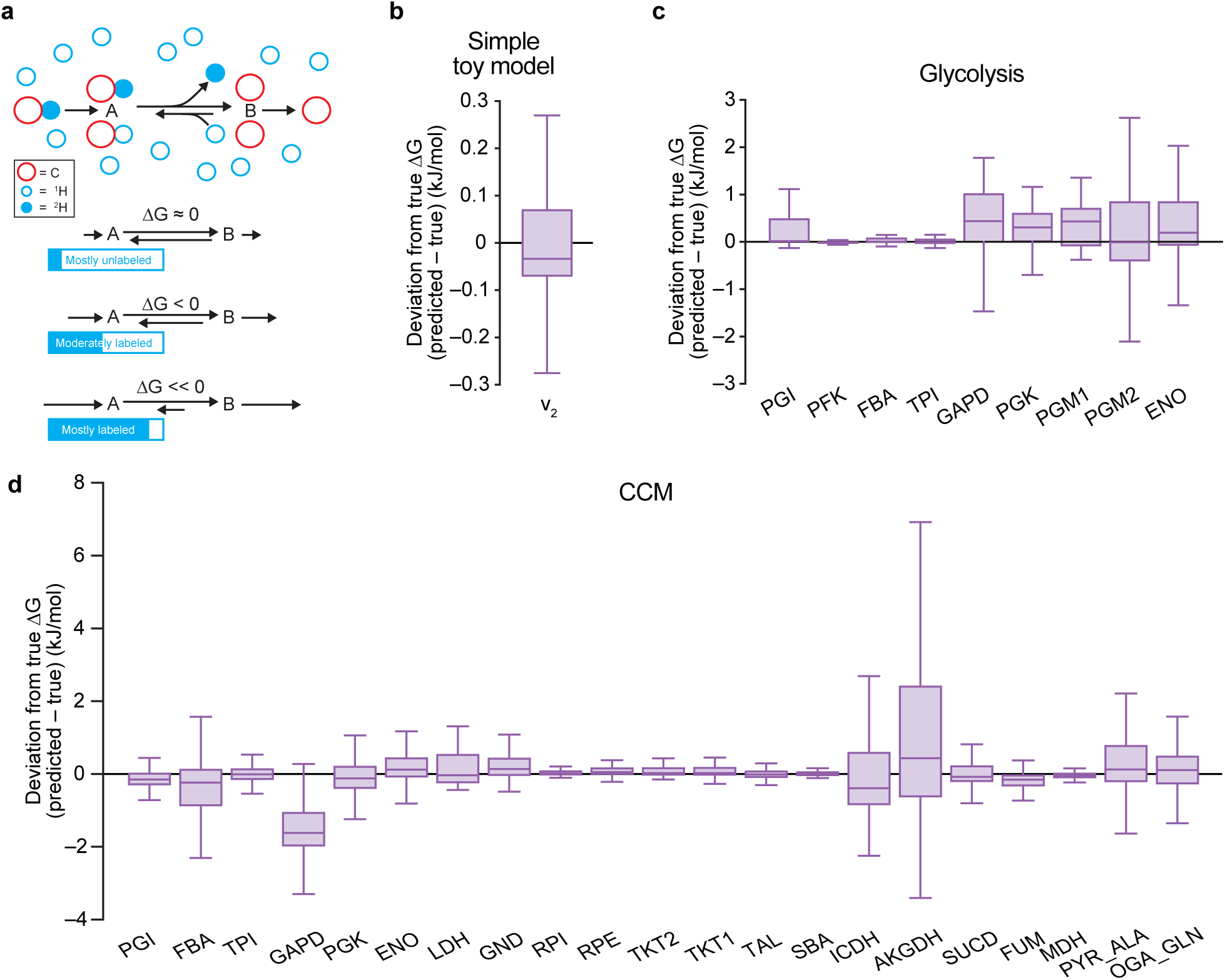
Deep learning of isotope labeling patterns reveals reaction free energy. **a,** Isotope labeling patterns reveal reaction reversibility and Gibbs free energy of reaction (ΔG). The blue circles represent hydrogen (empty) and deuterium (filled). Once deuterium is lost in the forward reaction, the reverse reaction picks up hydrogen. **b-d,** ML-Flux was used to assess the accuracy of ΔG prediction in (**b**) the simple toy model, (**c**) glycolysis, and (**d**) central carbon metabolism. The distribution of ΔG prediction errors was determined by sampling ranges of ΔG that were near equilibrium, highly reversible, or reversible (see **Methods**). Each box shows the three quartiles and whiskers extend to the minimum and maximum values within 1.5-fold of the interquartile range (n=300 for each flux for all models).

We tested the ability of ML-Flux to accurately predict ΔG across our five metabolic models. In the simple toy model, which has only one reversible reaction, the majority of ΔG prediction was within 0.1 kJ/mol of the true value (**Fig.5b**). We predicted ΔG across glycolytic reactions from ^2^H and ^13^C labeling patterns, where most errors fell within 1 kJ/mol (**Fig. 5c**). The CCM model accurately predicted ΔG for 21 reactions (**Fig. 5d**). The thermodynamic driving forces in nearly all central carbon metabolism reactions were predicted within 1 kJ/mol of their true value. The reaction catalyzed by α-ketoglutarate dehydrogenase (AKGDH) displayed larger errors in ΔG predictions but still within an interquartile range of –1 to 2 kJ/mol, a small relative error considering AKGDH is strongly forward-driven with ΔG<<0 kJ/mol. Overall, ML-Flux predicted ΔG accurately despite the incomplete isotope pattern input reflective of real-world experiments and the large cumulative ΔG across metabolic pathways (|ΔG|=∼30-100 kJ/mol)^46,47^. Since reaction thermodynamics provides a guiding principle for systems-level control of metabolism^4,37^, ML-Flux showed the potential to translate flux quantitation to actionable information for engineering metabolism.

### ML-Flux outperforms iterative solver-based MFA

We benchmarked ML-Flux against a leading MFA software package in terms of accuracy and speed of flux prediction (**Extended Data Fig. 8a**). Present MFA software searches the flux space for fluxes that simulate isotope labeling patterns with the least-squares discrepancy between simulated and measured isotope patterns. Over 90% of the time, ML-Flux computed flux more accurately than a leading MFA software (**Fig. 6a,b** and **Extended Data. Fig. 8b**). More accurate flux prediction by ML-Flux led to lower sum of squared residuals (SSR) of isotope labeling patterns compared with MFA software (**Extended Data Fig. 8c-e**). ML-Flux performance was consistent regardless of the availability of transport flux measurements (e.g., lactate secretion rates) that present MFA software relies on for accurate flux predictions (**Extended Data Fig. 8b,c**). In addition to higher accuracy, ML-Flux computed fluxes faster than MFA software did across all tested models (**Fig. 6c**) and combinations of isotope tracer experiments (**Fig. 6d**). With increasing iterations of least-squares optimization, present MFA software may equal or surpass the accuracy of ML-Flux but at the cost of time and computing power. Nevertheless, ML-Flux demonstrated its utility with orders of magnitude superior accuracy and speed.

**Figure 6.**
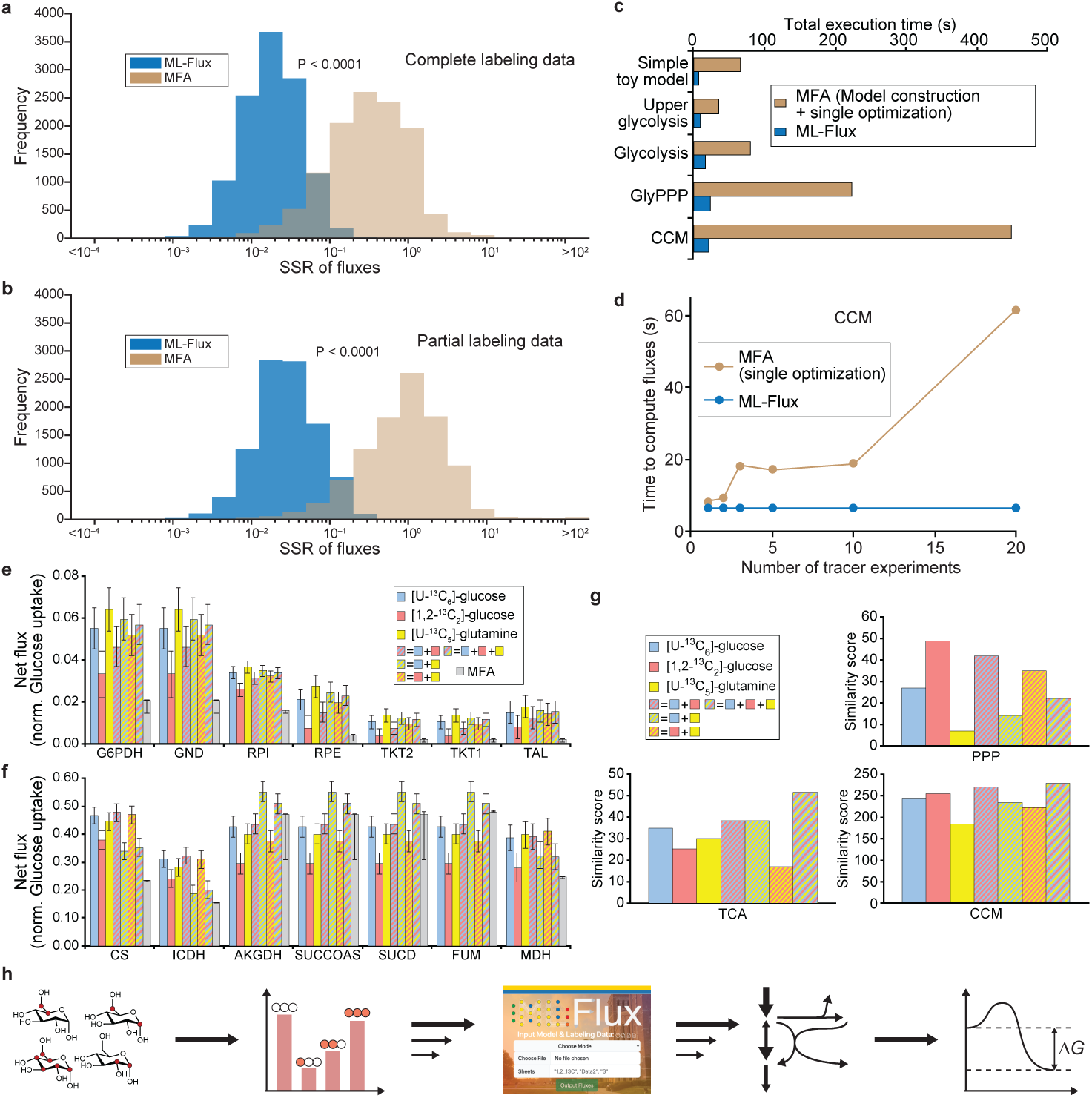
ML-Flux’s improved accuracy and speed leads to real-world applicability. **a** and **b,** Fluxes were predicted with either ML-Flux or a leading iterative solver-based MFA software using an input of either complete (**a**) or partial (**b**) isotope labeling patterns from the testing dataset. The sum of squared residuals (SSR) between the predicted and true fluxes were compared between the two methods. Reported statistics are from a logarithmically spaced two-tailed t-test. **c,** Total execution times for model construction and flux prediction were measured for ML-Flux versus least-squares MFA. **d,** For the CCM model, the time to compute fluxes was measured as a function of the number of parallel isotope tracer experiments. **e,** Net fluxes in the pentose phosphate pathway (PPP) were determined using ML-Flux with isotope pattern measurements from [U-^13^C_6_]-glucose, [1,2-^13^C_2_]-glucose, and/or [U-^13^C_5_]-glutamine tracing. ML-Flux results were compared to iterative solver-based MFA results from all three tracers. **f**, Net fluxes in the TCA cycle were predicted using the same tracer configurations and compared to results from MFA. Error bars in **e** and **f** reflect the error propagated from the standard error of individual flux predictions and of replicate predictions (n=3). For MFA, error bars represent the lower and upper bound of the flux value based on a 95% confidence interval analysis. **g,** Similarity scores were computed by ranking ML-Flux results from different tracer combinations by their proximity to the MFA-predicted values for the PPP, the TCA cycle, and the CCM networks. **h,** ML-Flux was deployed online for accurate and rapid determination of metabolic flux and free energy.

### ML-Flux turns real-world metabolomics data into accurate fluxes

We evaluated the performance of ML-Flux using real-world metabolomics data from mammalian cells cultured on [U-^13^C_6_]-glucose, [1,2-^13^C_2_]-glucose, or [U-^13^C_5_]-glutamine^46^. There is no ground truth of measured central carbon metabolic fluxes to confirm a flux prediction from real data. Instead, we compared ML-Flux predictions to those of MFA to see how the two approaches, each having their own accuracies, differ. We prepared seven subsets of isotope pattern data (from combinations of one, two, or three tracers) as input for ML-Flux and obtained fluxes through central carbon metabolism. In the PPP, ML-Flux predictions intervals from tracing [1,2-^13^C_2_]-glucose were within 0.01 flux units of MFA flux predictions using isotope labeling data from all three tracers (**Fig. 6e**). In the tricarboxylic acid (TCA) cycle, isotope labeling data from all three tracers produced fluxes within the confidence intervals within 0.05 flux units of most MFA fluxes (**Fig. 6f**). We determined similarity scores for the tracer combinations based on the closeness of individual net and exchange flux predictions between ML-Flux and MFA (**Fig. 6g** and **Extended Data Fig. 9a,b**). Benchmarking ML-Flux against MFA demonstrated how optimization for accurate flux predictions (as in ML-Flux) versus minimized residuals of isotope labeling pattern data (as in MFA) yields slightly different fluxes using real-world isotope labeling pattern data. Nonetheless, ML-Flux demonstrated its real-world applicability in flux determination.

Compared to MFA software using iterative solvers, ML-Flux requires little computational resources. Instead of iterative simulation of metabolite isotope labeling patterns and gradient descent algorithms, ML-Flux uses pre-trained models with most of the computational demand frontloaded in the form of neural network training. Low computational demand enabled the online deployment of ML-Flux, engendering the first open-source, open-access web tool for metabolic flux and free energy quantitation: metabolicflux.org. The ML-Flux web tool offers fast and accurate computation of fluxes and free energies with standard errors for individual isotope tracing experiments as well as standard deviations of multiple predictions from replicate experiments. The ML-Flux software package and web tool are regularly updated to incorporate new metabolic models and increase the speed and accuracy of flux quantitation. Our metabolic models and machine learning models, along with documentation, example data, and public repositories, constitute a growing knowledgebase of metabolic fluxes and free energies (**Fig. 6h**).

## Discussion

The advancement of analytical and computational tools has increased the sensitivity and coverage of nucleic acid, protein, and metabolite measurement, propelling genomics, transcriptomics, proteomics, and metabolomics^7,48,49^. However, fluxomics has lagged behind. The rates of metabolic reactions are determined by the interaction of enzymes and metabolites, which serve as substrates, products, and effectors. Thus, fluxomics plays an indispensable role in understanding biological systems in action and provides the missing link in integrative omics^50^. In this work, we advanced fluxomics and contributed to integrative omics by innovating two-stage machine learning.

Metabolic flux quantitation hinges on the relationship between isotope labeling patterns imprinted on metabolites and the underlying metabolic fluxes. In simple metabolic networks, the relationship appears algebraically straightforward^19^. Increasing complexity of metabolic networks renders these relationships nearly incomprehensible to human cognition^51,52^. By training machine learning models with copious isotope pattern-flux pairs sampled from a physiologically relevant flux space, ML-Flux captured inconspicuous isotope pattern-to-flux relationships. Accurate exchange flux quantitation by ML-Flux additionally led to resolving ΔG across central carbon metabolism. Connecting isotope patterns to not only net fluxes but also free energies expands our ability to target flux controlling steps in pathways^4,37^.

A challenge in training ML models for flux prediction was the limited availability of flux data in the literature, which predominantly originate from iterative solver-based MFA. Limited availability of ground truth flux data hinders training ML models^53^. To overcome this challenge, we generated expansive isotope tracing simulations across central carbon metabolism. Our simulations employed computationally efficient elementary metabolite units (EMU)^15^ to pair metabolic fluxes to corresponding isotope labeling patterns. Therefore, ML-Flux effectively frontloaded computational tasks and obviates extensive runtime computation unlike MFA, which performs computationally intensive least-squares algorithms at runtime. As a result, ML-Flux framework gave rise to near instantaneous flux quantitation online on demand for the first time.

Another benefit of ML-Flux was its flexibility in accommodating various choices of isotope tracers and metabolite input unlike conventional ANN models, which require fixed input sizes^54,55^. We accomplished this feat by training our ANN models using a comprehensive set of commercially available isotope tracers and incorporating PCNN for imputation of metabolite labeling patterns that are often not or cannot be measured. Although the imputation step adds the possibility for potential erroneous predictions, thorough training of the two-stage PCNN-ANN framework remained robust for nearly all conceivable isotope tracing in central carbon metabolism. Learning from multiple isotope tracers also led to the discovery of informative tracing experiments. While parallel tracing experiments with [1,2-^13^C_2_]-glucose and [U-^13^C_5_]-glutamine have been a long-established method for quantifying central carbon metabolism^43,44^, ML-Flux revealed that dual [4-^13^C_1_]-glucose and [U-^13^C_5_]-glutamine tracing can resolve fluxes equally well or better. Thus, machine learning helps researchers design tracing strategies for optimal flux determination.

ML-Flux and iterative solver-based MFA are different in two ways. The former focuses on accurate prediction of fluxes, whereas the latter focuses on accurate simulation of isotope patterns, causing it to be susceptible to multiple flux solutions (i.e., local optima) that yield similar isotope patterns within tolerance. The two approaches to flux quantitation also differ in how they propagate knowledge. ML-Flux preserves label-to-flux relationships that are learned permanently, whereas MFA does not store knowledge for long-term learning. Thus, the value of ML-Flux is that the capital effort of metabolic model construction and simulation is not expended in a single prediction. Future development of more sophisticated ML architecture will broaden and reinforce the utility of ML-Flux.

ML-Flux serves both experts and newcomers to flux analysis by playing dual roles of streamlining accurate flux quantitation and producing a sound starting point for a potential refinement by iterative solvers. The implementation of ML-Flux as a web tool with curated metabolic models and deep learning models establishes a resourceful and accessible knowledgebase for studying the metabolism of various organisms from microbes to humans. Thus, the upshot of ML-Flux is the democratization of metabolic fluxes for the acceleration of biotechnology and medicine.

## Methods

### Generation of metabolic models

Five metabolic models were designed for the simulation of isotope labeling patterns and training for ML-Flux: 1) a simplified upper glycolysis and the pentose phosphate pathway, 2) upper glycolysis 3) full glycolysis for the determination of reaction free energies, 4) glycolysis and the pentose phosphate pathway specialized, and 5) central carbon metabolism (with cytosolic oxaloacetate transport into the mitochondria as malate lumped with ATP citrate lyase). The first four models contained one metabolite input (A’ for model 1 or glucose for models 2-4). The central carbon metabolism model contained glucose and glutamine as input metabolites. All intracellular reactions in each metabolic model contained net and exchange fluxes to reflect that enzyme-catalyzed reactions are reversible with forward and reverse components. Physiologically relevant direction and reversibility were defined as linear equality and inequality constraints. Metabolic models were constructed and coded in an Extensible Markup Language (XML) format for simple and standardized archiving and sharing. Reactions considered in each model are detailed in **Supplementary Tables 2-5**.

### Analytical solutions to the simple toy metabolic model

Analytical solutions to fluxes as a function of isotope patterns were determined in the simple toy metabolic model by applying metabolic steady-state mass balance to all metabolite levels as well as isotopic steady-state mass balance to all isotopologues. Metabolic steady state is defined as:

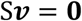

S is the stoichiometric matrix with rows and columns representing metabolites and reactions. ***v*** is the vector of fluxes. These balances determine linearly dependent relationships between fluxes:

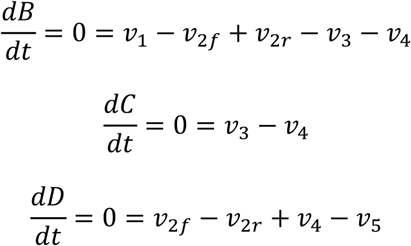

Isotopologue mass balance further relates MID measurements to fluxes. Using a [1-^13^C_1_]-A tracer, balances on B_M+1_ and D_M+1_ were used to obtain fluxes for v_3_ and v_2r_.

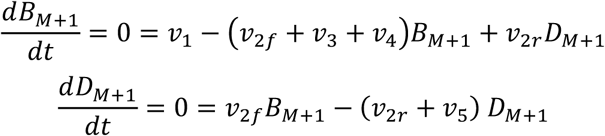

These equations were rearranged in terms of v_1_,v_3_, and v_2r_. Normalizing all fluxes to that of a reference reaction (i.e., v_1_=1 in this example), yields the solutions to v_3_ and v_2r_. The solved fluxes can be reinputted into metabolite balances of the entire model to obtain all fluxes in the system (**Fig. 1f**).

### Sampling of metabolic fluxes for training ANN models

Fluxes for simulation of metabolite isotope labeling patterns were sampled within a physiologically relevant flux space (**Supplementary Tables 2-5**). At a metabolic steady state, the net fluxes of the system reside in the null space of S. Free net fluxes were found from its reduced row echelon form using the rank-nullity theorem, and the multiplication of the basis of the null space by free net fluxes returned the full set of net fluxes. All exchange fluxes were free variables. Free net fluxes were sampled using a linearly uniform random distribution. Exchange fluxes were generated using either a linearly uniform, logarithmically uniform, or logarithmically normal random distribution. To generate a training set containing fluxes that simulate a near uniform distribution of isotope patterns in the simple toy model, distance-weighted sampling was employed (**Extended Data Fig. 1c,d**). Each newly sampled fluxes simulated isotope patterns, which were accepted into the final dataset based on a probability proportional to the average Euclidean distance (D) of the 10 nearest isotope patterns already in the dataset. The probability was assigned as D/D_max_ where D_max_ is the maximum distance between any two points in the dataset. The final training datasets used for the simple toy, upper glycolysis, glycolysis, and GlyPPP models were sampled from a log-uniform distribution. For the simple toy, upper glycolysis, and full glycolysis models, 100,000 flux distributions were generated to train ML models.

For the GlyPPP and CCM model, we employed an artificial centering hit-and-run algorithm modified from the COBRA Toolbox to generate a sample of 1,000,000 flux sets for both models that uniformly cover the feasible flux space^9,56^. However, since the resultant sample set does not necessarily generate uniform distributions for individual fluxes, potential biases in the training data may persist. To reduce this bias in the CCM model, we conducted rejection sampling for six fluxes through LDH, PDH, PPCK, ME, MDH, AKGDH to mitigate overrepresentation of any flux state. The final CCM dataset was reduced from 1,000,000 to 117,077 flux distributions (**Extended Data Fig. 10**), a smaller but higher quality dataset to train from.

### Simulation of isotope labeling patterns

For each flux set, metabolite isotope labeling patterns were simulated starting from various isotope tracers (**Supplementary Table 1**). For the simple toy model, all possible tracer forms of metabolite ‘A’ were used for simulation except for the trivial set (i.e., fully labeled or fully unlabeled). For upper and full glycolysis models, [1,2-^13^C_2_]-glucose and [5-^2^H_1_]-glucose tracers were used. In the GlyPPP model, 24 commercially available ^13^C-glucose tracers were used. Isotope labeling patterns were simulated following the EMU approach^15^. In the CCM model, the top 10 most used ^13^C-glucose tracers according to Google Scholar were simulated with a combination of either unlabeled or [U-^13^C_5_]-glutamine for a total of 20 unique tracer combinations. Models in an XML format^29^ with atom mapping information were processed through MATLAB scripts to reconstruct them as a stoichiometric model and EMU networks. For isotope pattern simulation in the CCM model, we took into account symmetry of succinate and fumarate^15,57^. Using the sampled free net and exchange fluxes, the EMU framework was employed to find simplified isotopologue conversion networks and simulate mass isotopomer distributions based on atom mapping across reactions. The resulting pairs of reaction fluxes and metabolite isotope labeling patterns were stored in .dat files. Files were formatted such that each row contained isotope patterns from a single flux set, ordered by metabolite, the isotope tracer used, and then mass isotopomer fractions.

### Principal component analysis of the simple toy metabolic model

Simulated isotope patterns from the simple toy metabolic model were subjected to principal component analysis (PCA). To generate principal components directly related to metabolite isotope labeling patterns, and since fractions of mass isotopomers always fall in the range of [0,1], PCA input features were uncentered and unscaled. Since the sum of all mass isotopomer fractions for a metabolite is always one, uniformly unlabeled isotopomer fraction was excluded from PCA, and only the fractions of independent mass isotopomers harboring heavy isotopes were used in PCA.

### Flux prediction by artificial neural networks

Simulated flux-to-label simulations were used to train five unique fully connected feedforward ANNs. The input layer for each of these ANNs was the complete isotope labeling information of every metabolite within the metabolic model for every isotope tracer used. The output layer was the free fluxes of the metabolic model. Training under the set of free fluxes ensured that the minimal linearly independent output nodes were used, and the resulting full flux set was always mass balanced.

Isotope pattern-flux data pairs were split into training, validation, and testing at a ratio of 0.8:0.1:0.1. ANN training scripts were written in Python using the Keras library. Input to ANNs were metabolite isotope labeling patterns, and the output was transformed free fluxes, which were later reverted to the actual values by inverse functions. Each neural network contained one input layer for MIDs, five hidden layers, and one output layer for free fluxes (**Supplementary Table 9**). The number of nodes within each hidden layer was chosen empirically in each ANN depending on the complexity of the metabolic models. For the first four metabolic models, fluxes were transformed under a piecewise logistic or logarithmic function to minimize the effect of very large fluxes. The corresponding ANNs were optimized using a mean absolute error loss function. For the CCM model, a custom loss function that calculated the weighted mean squared errors across all fluxes in the metabolic model was used. Weights were assigned as the reciprocal of the upper limits of individual flux constraints. For example, a flux that was constrained from 0-0.2 during flux sampling would have a weight of 5. The ANN model for CCM model was optimized under this loss function twice: first with all ANN parameters trainable, followed by tuning parameters other than those related to transaldolase flux (the free flux determinant of PPP fluxes) were frozen. Training was conducted on the Purdue Anvil cluster, the UCLA Hoffman2 cluster, and on local workstations. The standard errors of fluxes were derived from half the range of the center 68% of the distributions of prediction errors from testing data (**Supplementary Tables 6 and 7**).

### Imputation of isotope patterns by partial convolutional neural networks

Simulated isotope labeling patterns were masked to emulate the proportion of incomplete information due to detecting a subset of metabolites or performing a subset of isotope tracer experiments (**Supplementary Table 10**). Masks randomly removed isotope labeling patterns of various fractions of metabolites from all but one to a few isotope tracers. The masked isotope patterns were used for training PCNN inpainting models using the Keras library (**Supplementary Tables 11-15**). The isotope labeling pattern data were reshaped into a rectangular matrix and overlaid on the center of a square matrix. Extra space was padded with ones and not masked. The sizes of the square matrix ranged from 16×16 to 80×80 depending on metabolic networks. The incomplete-complete matrix pairs were split into training, validation, and testing at a ratio of 0.8:0.1:0.1. Mean absolute error or mean squared error was used for the loss function. PCNN-based inpainting training was conducted using Nvidia A100 GPU nodes on the Anvil cluster.

The plausibility of inpainted MID matrices was quantified by the following equation (near-zero ΔN values indicate high plausibility of inpainted isotope labeling patterns):

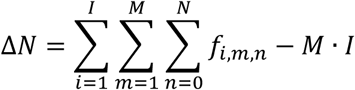

*i* represents an isotope tracer experiment simulated in the model, *m* represents a metabolite in the pathway model, *n* represents a mass isotopomer M+n, and f_i,m,n_ represents the fraction of M+n isotopomer of metabolite m given an isotope tracer *i*. *M* is the total number of metabolites in the metabolic model, *I* is the total number of isotope tracers used for training ML models, and *N* is the maximum number of atoms considered in atom mapping in a metabolite.

### Calculation of ΔG in upper glycolysis and glycolysis

Gibbs free energy of reaction was calculated using net and exchange fluxes:

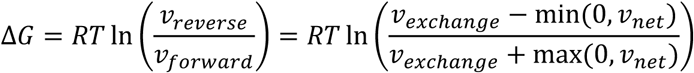

R is the universal gas constant, and T is temperature in Kelvin. To compare between the ΔG values from testing data and ML-Flux predictions, test data for each flux was evenly sampled from three bins that represent physiological ranges: near equilibrium, –0.5 kJ/mol ≤ ΔG ≤ 0 kJ/mol; highly reversible, –1 kJ/mol ≤ ΔG < 0.5 kJ/mol; and reversible, ΔG < –1 kJ/mol. The errors of ΔG prediction for each reversible reaction was determined as half the range of the center 68% of the distributions of prediction errors from the sampled testing data.

### Goodness-of-fit determined by reduced χ^2^ of fluxes

Since the goal of flux analysis is to accurately predict fluxes from given isotope tracing experiments, we performed a statistical test to quantify the accuracy of flux prediction. To this end, a reduced χ^2^ test was employed using the variance-weighted summed squared residuals (VSSR) of fluxes.

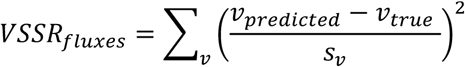

*v* represents the predicted flux and the true flux, and *s_v_* represents the standard deviation of the corresponding flux. We determined a critical VSSR based on a χ^2^ distribution with degrees of freedom equal to the number of free (independent) fluxes and significance level α=0.05. Predicted fluxes with a VSSR less than the critical value were deemed statistically acceptable.

On the other hand, iterative solver-based MFA searches for fluxes whose simulated isotope patterns best match that of the experimentally measured values by minimizing VSSR_MIDs_.

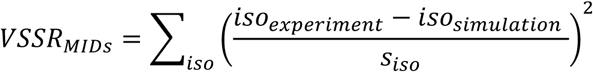

*iso* is the MID (i.e., isotope labeling patterns) from experiment or simulation, and *s_iso_* is the standard deviation of the measured MID. A flux set is deemed statistically acceptable if its VSSR of MIDs is lower than a critical VSSR determined by a χ^2^ distribution^58^.

The trained ML models were subjected to statistical analyses using a reserved testing dataset with known true fluxes. VSSR_fluxes_ values computed across a range of relative standard deviations of fluxes revealed the proportion of test predictions that pass the reduced χ^2^ test as a function of tolerance for errors (**Extended Data Fig. 4b**). The tolerances for accepting 95% of net and exchange fluxes were ∼0.1 and ∼0.68, respectively, which were comparable to or better than the errors given in confidence-interval analysis from iterative solver-based MFA^59^.

### Comparison of flux prediction performance with iterative solver-based MFA

Using the same testing data and the same metabolic model of central carbon metabolism, ML-Flux was compared with a leading iterative solver-based MFA. Two scenarios with complete and incomplete isotope pattern data were considered. For the former, the input encompassed the isotope labeling patterns of all metabolites from all 20 isotope tracer combinations. For the latter, a subset of isotope patterns chosen from 10-30 randomly selected metabolites from one or two isotope tracer experiments. Comparison of flux prediction accuracy were carried out with or without the inclusion of transport fluxes in the input to iterative solver-base MFA. ML-Flux and iterative solver-base MFA were compared in flux prediction accuracy and resolution (VSSR_fluxes_ and VSSR_MIDs_) as well as their speed.

### Similarity score between ML-Flux and iterative solver-based MFA

Fluxes in mammalian central carbon metabolism were determined using ML-Flux and experimental measurement of isotope labeling patterns from [U-^13^C_6_]-glucose, [1,2-^13^C_2_]-glucose, or [U-^13^C_5_]-glutamine tracing. Seven combinations of one, two, or three of the isotope tracers resulted in seven flux predictions, which were compared to the results from iterative solver-based MFA. A similarity score for each tracer combination was defined to rank how ML-Flux results resembled that of MFA.

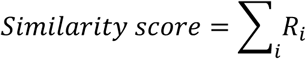

*R_i_* represents the rank (out of the seven tracers) of a tracer combination in its ability to predict to flux *i*. An *R_i_* of seven was assigned to the tracer combination resulting in fluxes that were closest to those of MFA, while an *R_i_* of one was assigned to the tracer combination resulting in fluxes that were farthest from those of MFA. Thus, a higher similarity score corresponded to a tracer combination that resulted in overall flux distributions that were more consistent with the MFA results.

### Online deployment of ML-Flux

All machine learning model architectures were stored in .json files with accompanying weights and biases in .h5 files. Reading these files for model prediction required minimal software, enabling light deployment onto a website using a Python backend and the Bootstrap CSS framework. With detailed documentation, examples, and template input files, ML-Flux was deployed at metabolicflux.org for accurate and rapid flux determination online.

## Acknowledgements

The authors thank the members of the Park Lab for constructive feedback. This work was supported by the National Institute of General Medical Sciences of the National Institutes of Health under Award Number R35GM143127 (J.O.P.), the National Center for Complementary and Integrative Health of the National Institutes of Health under Award Number R21AT012694 (J.O.P. and P.K.L.), the Department of Energy under Award Number DE-SC0024251 (J.O.P. and P.K.L.), the Center for Clean Technology Fellowship (R.C.L.), the National Science Foundation Research Traineeship in Integrated Urban Solutions for Food, Energy, and Water Management (NSF-INFEWS) under Award Number DGE-1735325 (G.N.), and the BioPACIFIC Materials Innovation Platform of the National Science Foundation under Award Number DMR-1933487 (R.C.L., G.N., and J.O.P.). This work used computational and storage services associated with the Hoffman2 Shared Cluster provided by UCLA Institute for Digital Research and Education’s Research Technology Group as well as the Advanced Cyberinfrastructure Coordination Ecosystem: Services & Support (ACCESS) computing resources (CHM230023). The content is solely the responsibility of the authors and does not necessarily represent the official views of the National Institutes of Health, the Department of Energy, or the National Science Foundation.

## Code and data availability

The code for simulating isotope labeling pattern data and training machine learning models is available on the GitHub public repository: https://github.com/richardlaw517/ML-Flux

---

**Extended Data Figure 1.**
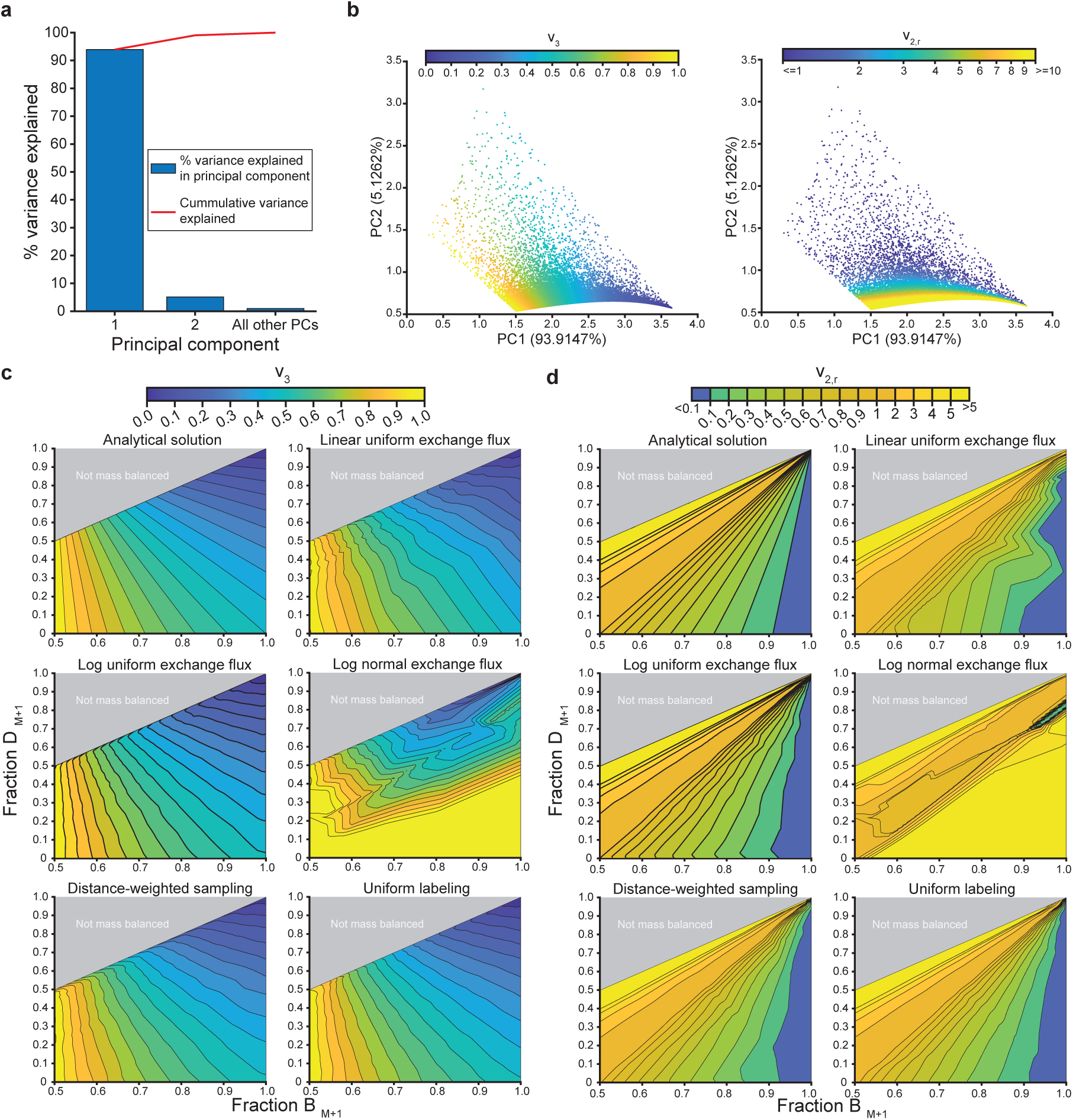
Different flux sampling methods to generate training datasets alter the outcome of ANN model training. **a,** Isotope labeling patterns from the simple toy metabolic model were used for principal component analysis, where each metabolite isotopomer from a given isotope tracer experiment was considered as a feature. The resulting dimension reduction revealed that 99% of the variance in the data could be captured in the first two principal components. **b,** Isotope labeling patterns were transformed onto the first and the second principal components with colors representing the magnitude of corresponding flux values for v_3_ (left) and v_2,r_ (right). **c** and **d,** Five different sampling approaches for generating flux-to-label simulations for training were tested and compared to the analytical solution (top left) for (**c**) v_3_ and (**d**) v_2,r_ (see **Methods**). Each panel shows the solution space for v_3_ or v_2,r_ using one of the sampling approaches.

**Extended Data Figure 2.**
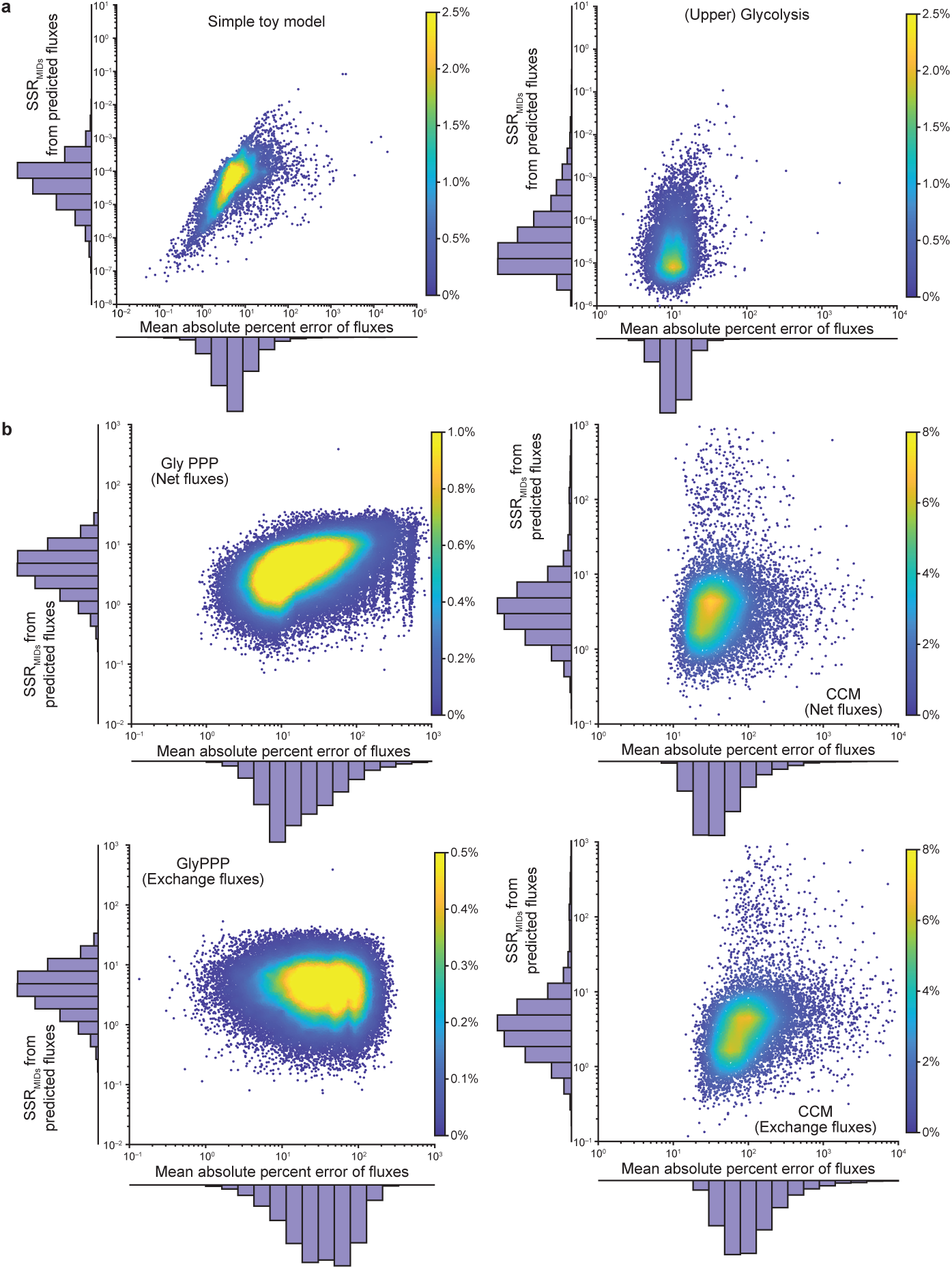
Accuracy of flux prediction and resolution of isotope labeling patterns characterize the quality of ANN flux predictions. **a,** For the simple toy and glycolysis metabolic models, ANN predictive capabilities were quantified by comparing fluxes and isotope patterns. The mean absolute percent error of predicted fluxes was calculated for each flux distribution in the testing data. Each predicted flux set was used to simulate isotope labeling data and calculate the sum of squared residuals between the isotope labeling patterns simulated from either the test or predicted fluxes. These two metrics were plotted (n=10,000 test flux-label sets). **b,** A similar analysis was conducted for the GlyPPP and CCM models with mean absolute percent errors calculated for net fluxes (top) or exchange fluxes (bottom) (n=100,000 and 11,708 test flux-label sets).

**Extended Data Figure 3.**
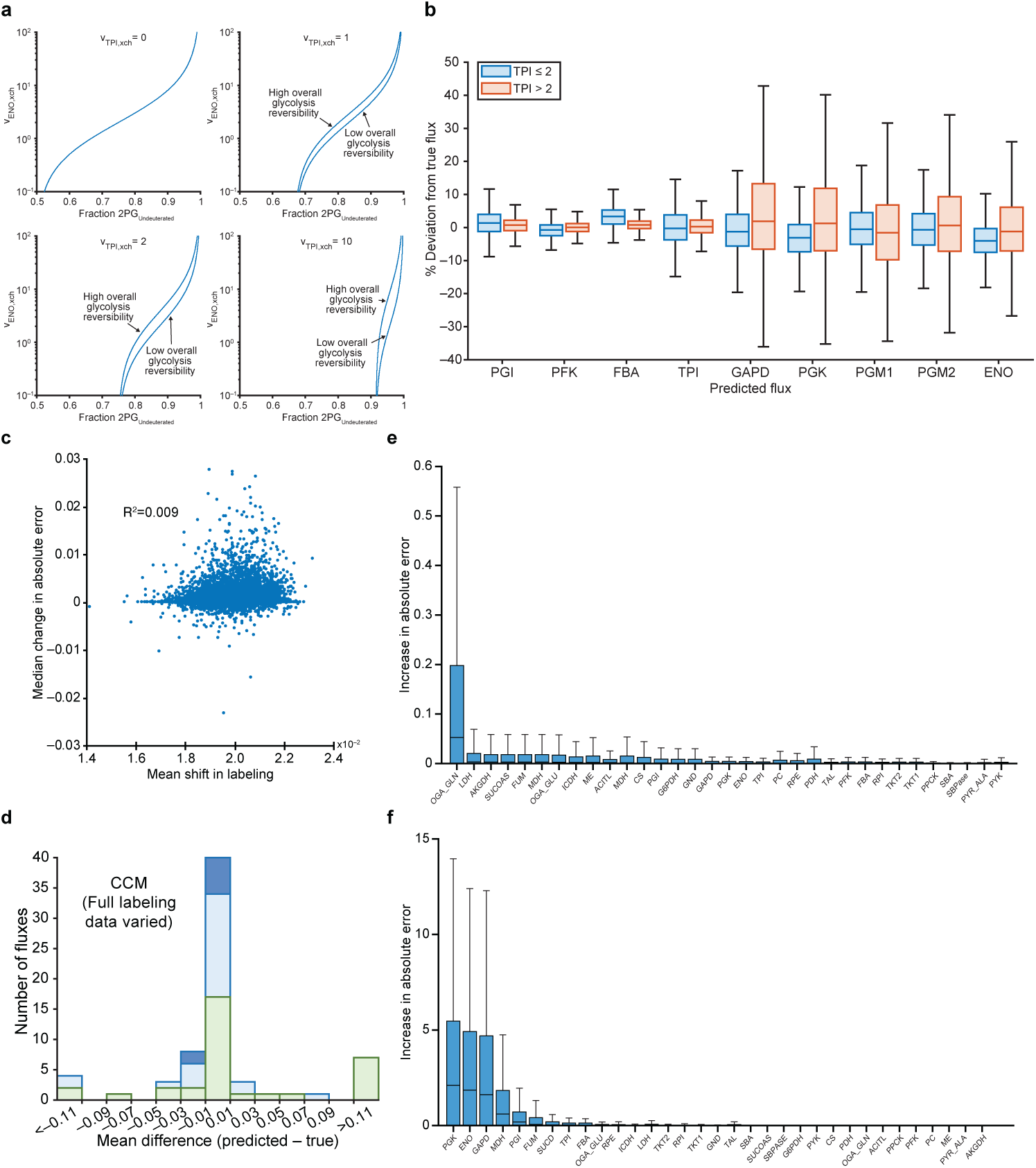
Fluxes and isotope patterns are sensitive to each other. **a,** The relationship between 2PG deuterium labeling and ENO exchange flux was determined by simulating isotope labeling data at logarithmically spaced intervals of ENO exchange fluxes with low (upper left) to high (lower right) TPI reversibility and all other reactions being either irreversible or highly reversible. With increasing magnitudes of TPI reversibility, the sensitivity of 2PG labeling to ENO exchange flux decreased. **b,** Test flux predictions in the full glycolysis model were separated by test fluxes with either low (≤2) or high (>2) TPI exchange flux. The resulting distribution of errors in flux predictions for these test sets were plotted, where each box shows the three quartiles, and whiskers extend to the minimum and maximum values within 1.5-fold of the interquartile range (n=430 for low TPI reversibility and n=9,570 for high TPI reversibility). **c,** Input MIDs for the CCM ANN model were varied by amounts consistent with typical instrumental error and inputted for ML-Flux prediction. The resulting change in error of flux prediction between the varied and non-varied inputs was calculated. **d,** The mean errors of fluxes in the CCM model were also calculated when using varied instead of exact isotope labeling data. **e** and **f,** The increase in error of individual net (**e**) and exchange (**f**) fluxes was measured between varied and exact isotope labeling inputs.

**Extended Data Figure 4.**
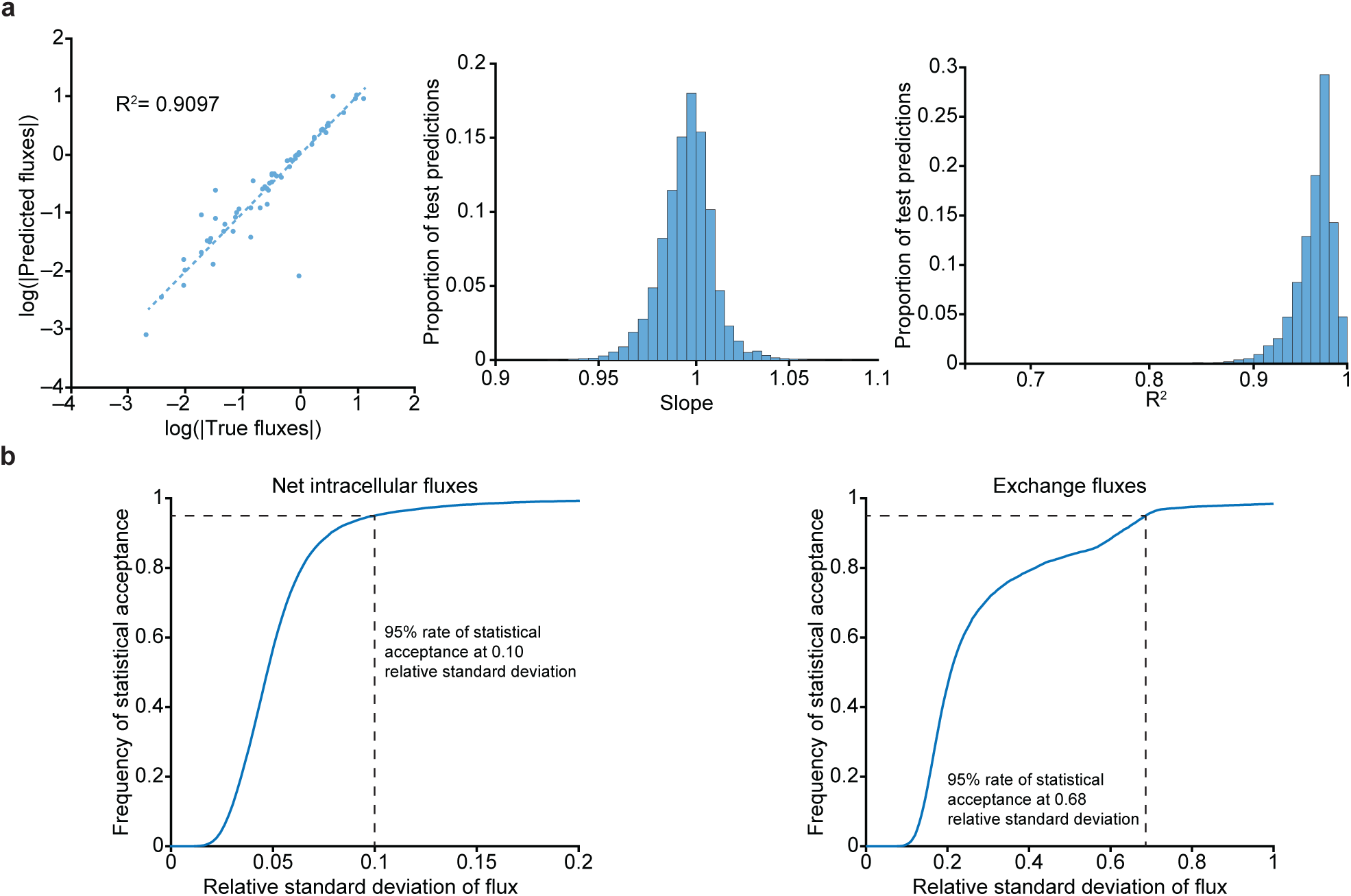
ML-Flux makes accurate and statistically acceptable flux predictions. **a,** Test flux predictions were compared to their true values by linear regression to determine the 1:1 relationship between predicted and true flux values (left). All test predictions are both mapped to a near 1:1 relationship (middle) and have a high correlation (right). **b,** Statistical validity of the trained ML models was confirmed using a reduced χ^2^ analysis. A reduced χ^2^ metric based on the summed residuals of flux predictions across a range of relative standard deviation was calculated. The frequency of the model giving statistically acceptable predictions was determined by proportion of predictions with a calculated χ^2^ was below the critical value determined by the number of free fluxes in the model (see **Methods**).

**Extended Data Figure 5.**
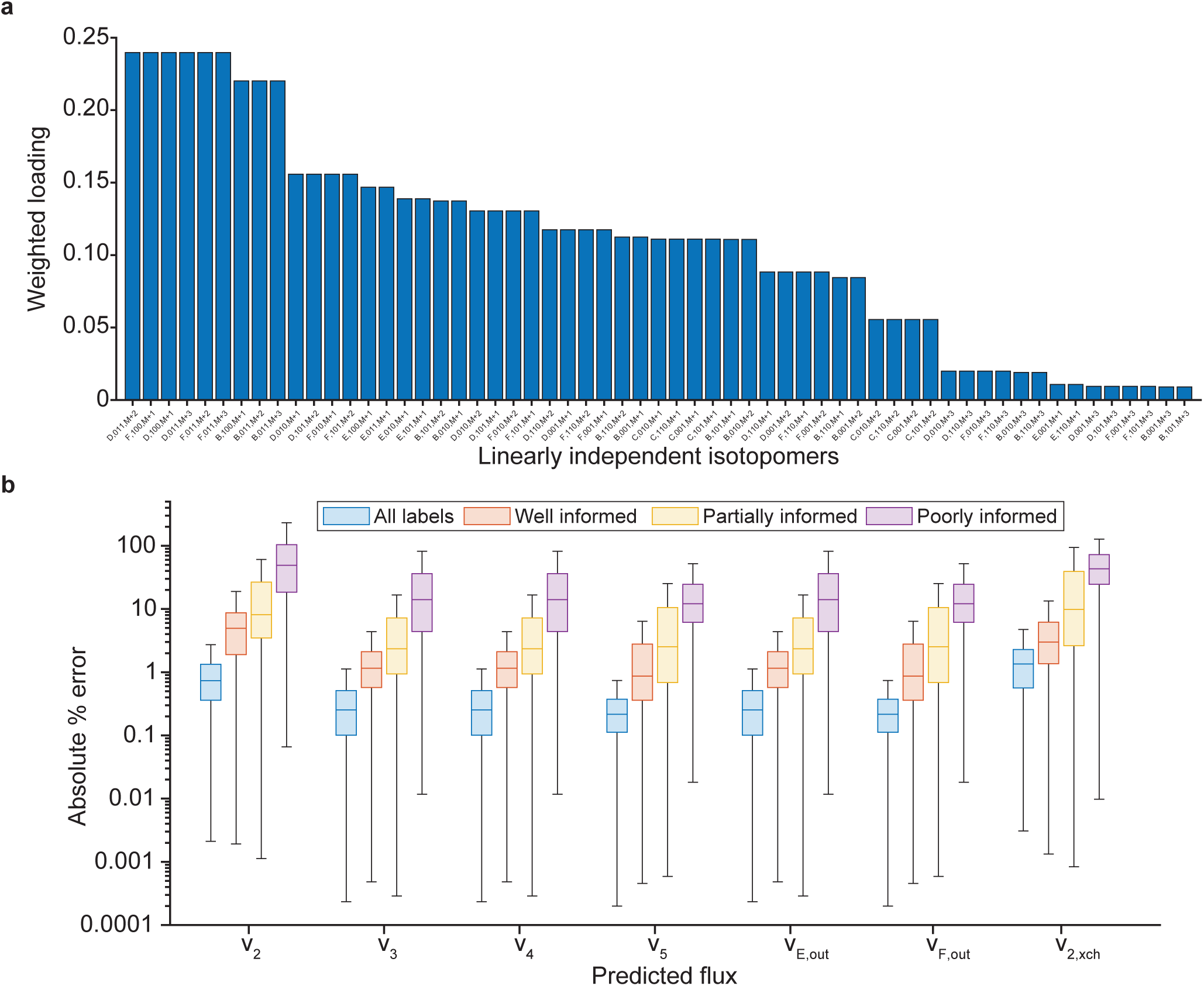
Accuracy of flux predictions depend on the measurement of key metabolite isotope labeling patterns. **a,** From PCA of the isotope labeling patterns in the simple toy model, the weighted sum of each linearly independent isotopomer’s loading for all principal components was calculated and sorted from high to low. Each bar across the x-axis describes a linearly independent isotopomer based on the metabolite, isotopologue of tracer A used, and isotopomer of the measured metabolite. For example, the first entry is the M+2 fraction of metabolite D from [011]-A tracing. **b,** Ranking of metabolite feature importance from PCA was used to sort test data from the simple toy metabolic model into four subgroups of fully informed (no isotope labeling data masked), well informed (both D and B isotope labeling unmasked), partially informed (either D or B unmasked) or poorly informed (neither D nor B isotope labeling unmasked). The resulting distribution of absolute percent errors in flux predictions were plotted. Each box shows the three quartiles, and whiskers extend to the minimum and maximum values within 1.5-fold of the interquartile range (n=100, 3,200, 5,800, and 1,000, respectively).

**Extended Data Figure 6.**
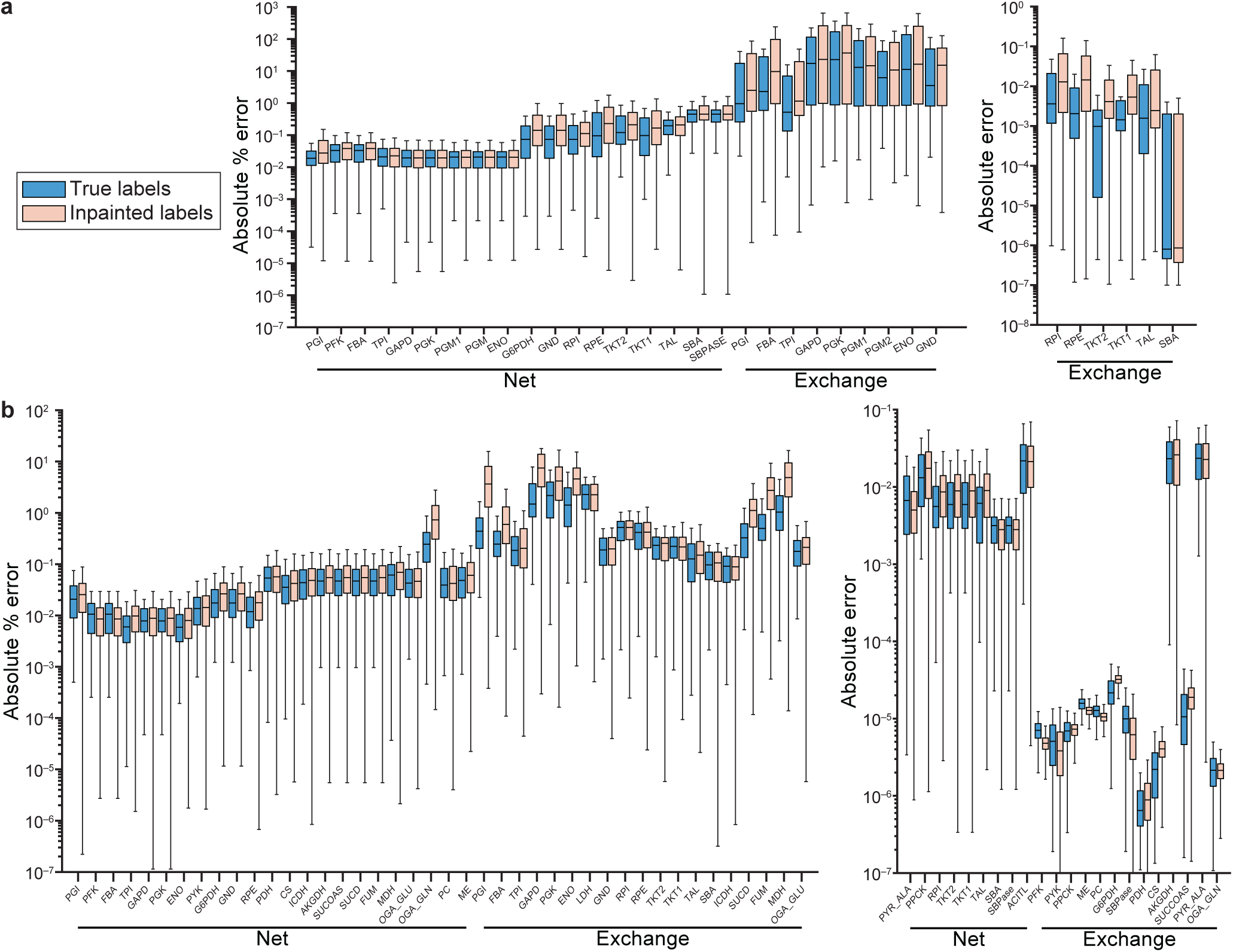
Integrated ML-Flux models predict net and exchange fluxes for larger metabolic models. **a-b,** Fluxes were predicted either from complete isotope labeling datasets or masked versions of the same datasets that undergo PCNN imputation. The distribution of their absolute errors was measured for the (**a**) GlyPPP and (**b**) CCM models. Fluxes with average values greater than 0.05 were assessed using absolute percent errors. Small deviations in fluxes with low true values have large relative errors but small impact on correctly capturing flux distributions. Thus, fluxes with average values less than 0.05 were assessed using absolute error. Each box shows the three quartiles, and whiskers extend to the minimum and maximum values within 1.5-fold of the interquartile range (n=100 for complete isotope labeling data, n=10,000 for masked isotope labeling data).

**Extended Data Figure 7.**
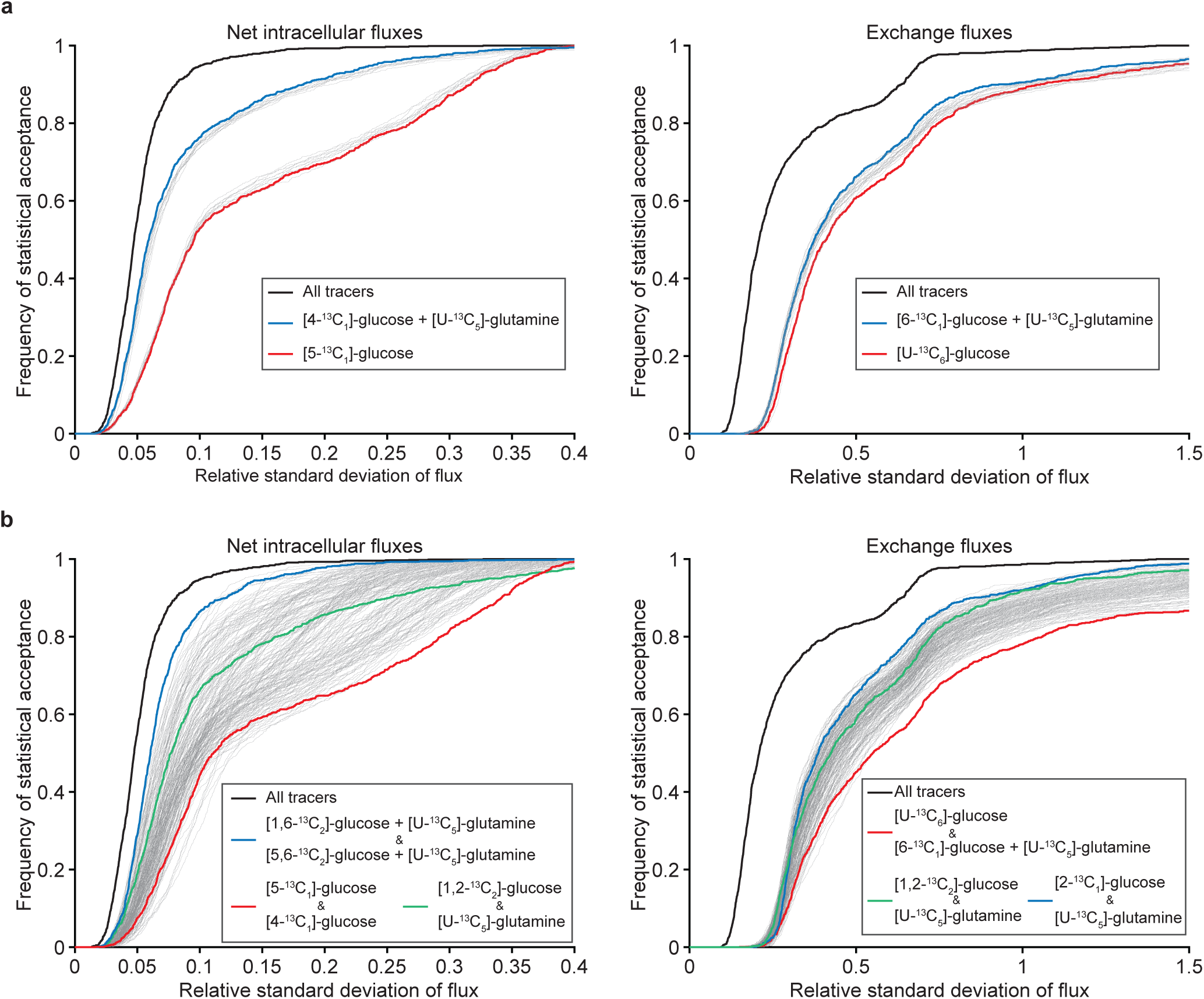
Select isotope tracers provide the most informative isotope labeling patterns for flux prediction. **a,** Metabolite isotope labeling patterns from single and dual isotope tracing experiments corresponding to 10 ^13^C-glucose tracers and 10 ^13^C-glucose-^13^C-glutamine tracer combinations were provided as input to ML-Flux for flux prediction (grey), which was compared to parallel tracing of all 20 tracer combinations (black). The frequency of passing the reduced χ^2^ statistical test of fit was plotted as a function of relative standard deviation (see **Methods**). The best (blue) and worst (red) performing tracer was then determined based on the relative standard deviation needed to achieve an 80% frequency of statistical acceptance. **b,** A similar analysis to **a** was conducted but for every combination of two parallel isotope tracing experiments (190 total), with the parallel tracing experiment of [1,2-^13^C_2_]-glucose and [U-^13^C_5_]-glutamine tracing highlighted in green.

**Extended Data Figure 8.**
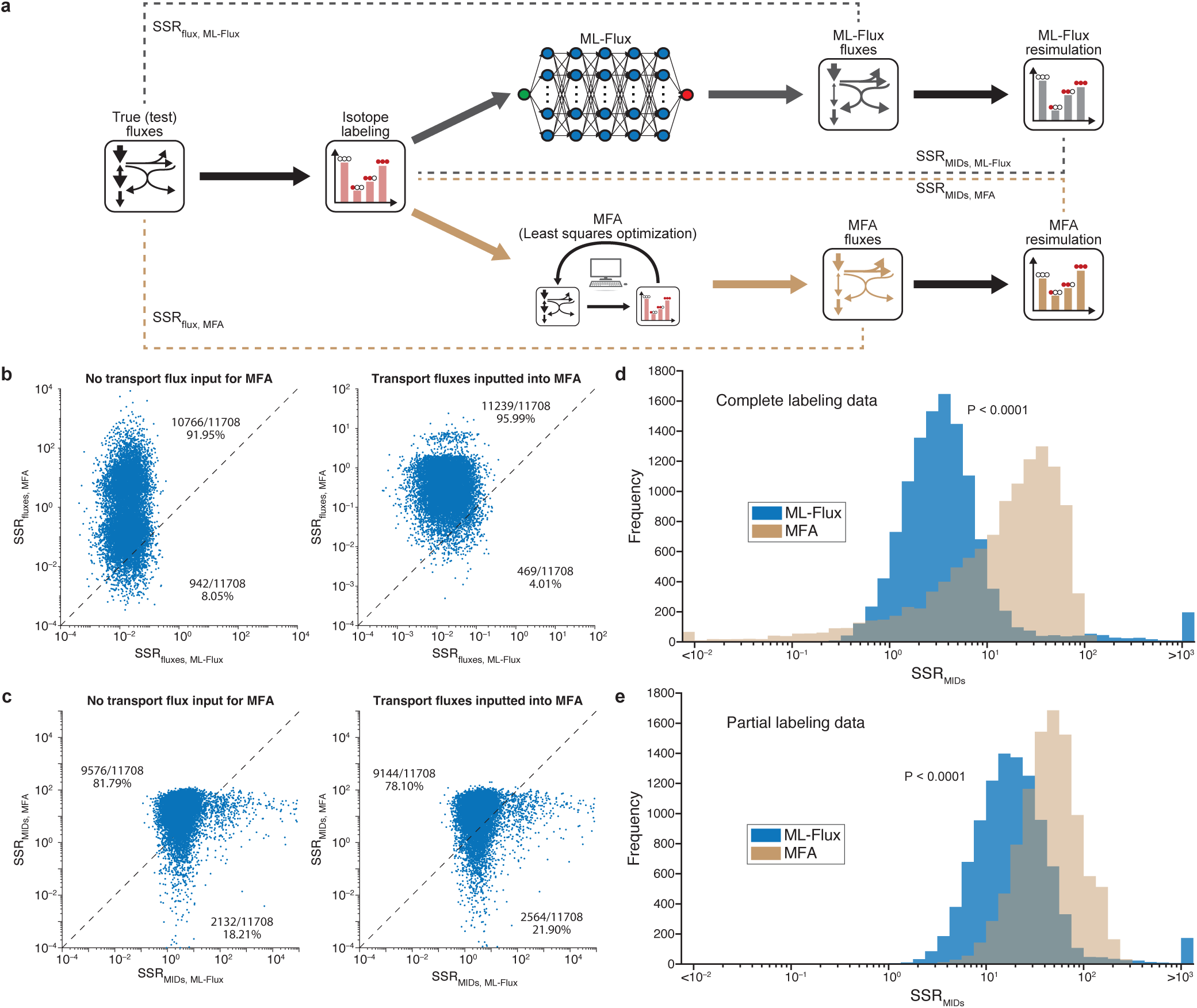
ML-Flux predicts metabolic fluxes at high accuracy. **a,** The quality of flux predictions was assessed using the summed squared residuals of fluxes or isotope patterns. The predicted fluxes were compared to the true flux values. The predicted fluxes were used to simulate isotope labeling patterns, which were compared to the input isotope labeling patterns. **b,** Comparison of the sum squared residuals of flux predictions was made between ML-Flux and MFA. In MFA, predictions were made without (left) or with (right) input of measured transport fluxes, while ML-Flux had no transport fluxes input in either case. **c,** A similar comparison was made using the sum squared residuals of MIDs simulated from the predicted flux values. **d-e,** The distributions of SSRs of MIDs for the test predictions were plotted for cases in which (**e**) the full isotope patterns or (**e**) variable-size isotope patterns were input into flux determination. Reported statistics are from a logarithmically spaced two-tailed t-test.

**Extended Data Figure 9.**
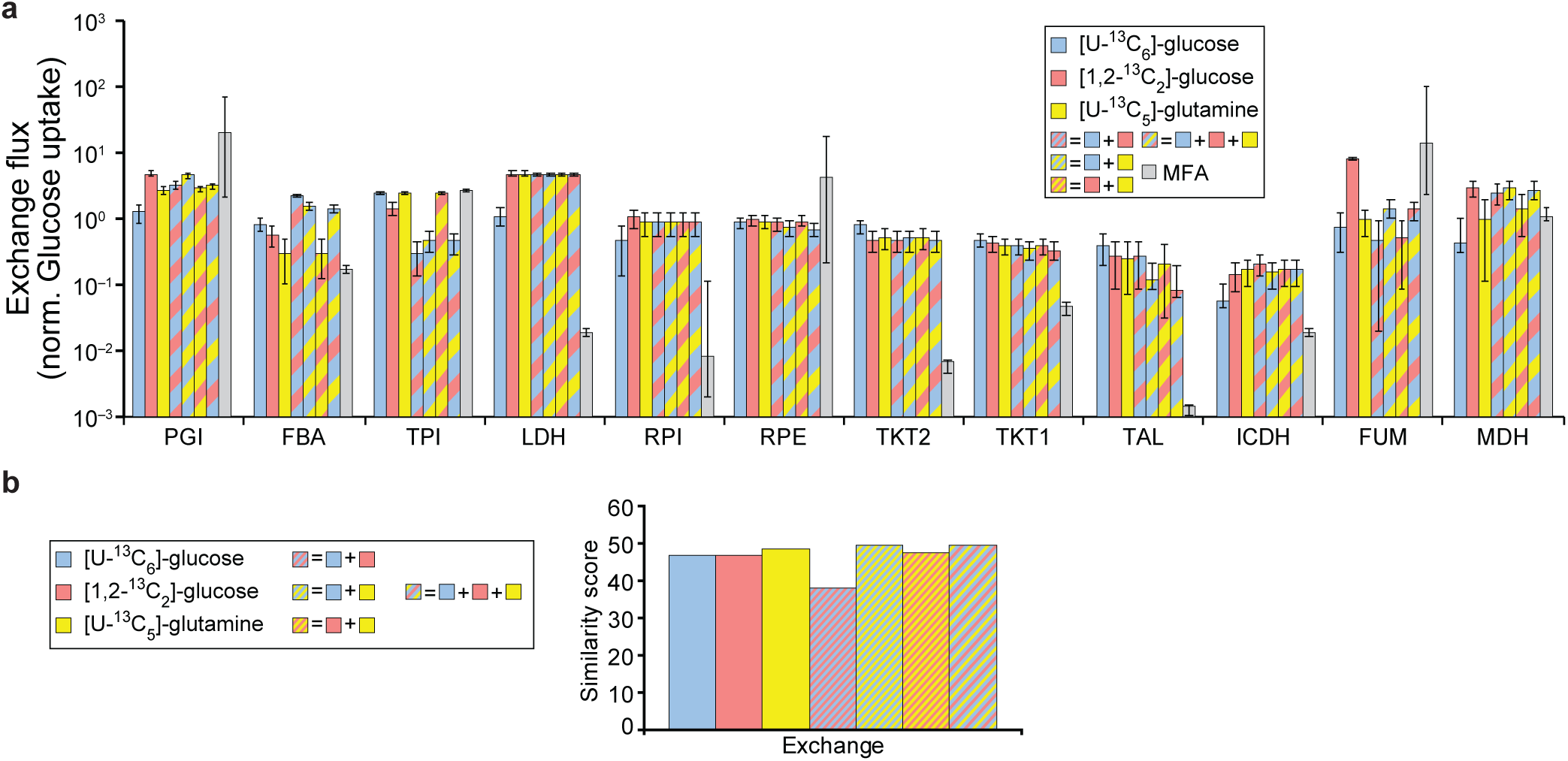
ML-Flux predicts exchange fluxes that are comparable to MFA results. **a,** Exchange fluxes were predicted using ML-Flux and various combinations of [U-^13^C_6_]-glucose, [1,2-^13^C_2_]-glucose, and [U-^13^C_5_]-glutamine tracers. Predicted fluxes were compared to those of MFA fitting all three tracing datasets simultaneously. Error bars reflect the error propagated from the standard error of individual flux predictions and of replicate predictions (n=3). For MFA, error bars represent the lower and upper bound of the flux value based on a 95% confidence interval analysis. **b,** The similarity scores were computed by ranking flux predictions from different tracer combinations by their proximity to the MFA predicted values for the exchange fluxes.

**Extended Data Figure 10.**
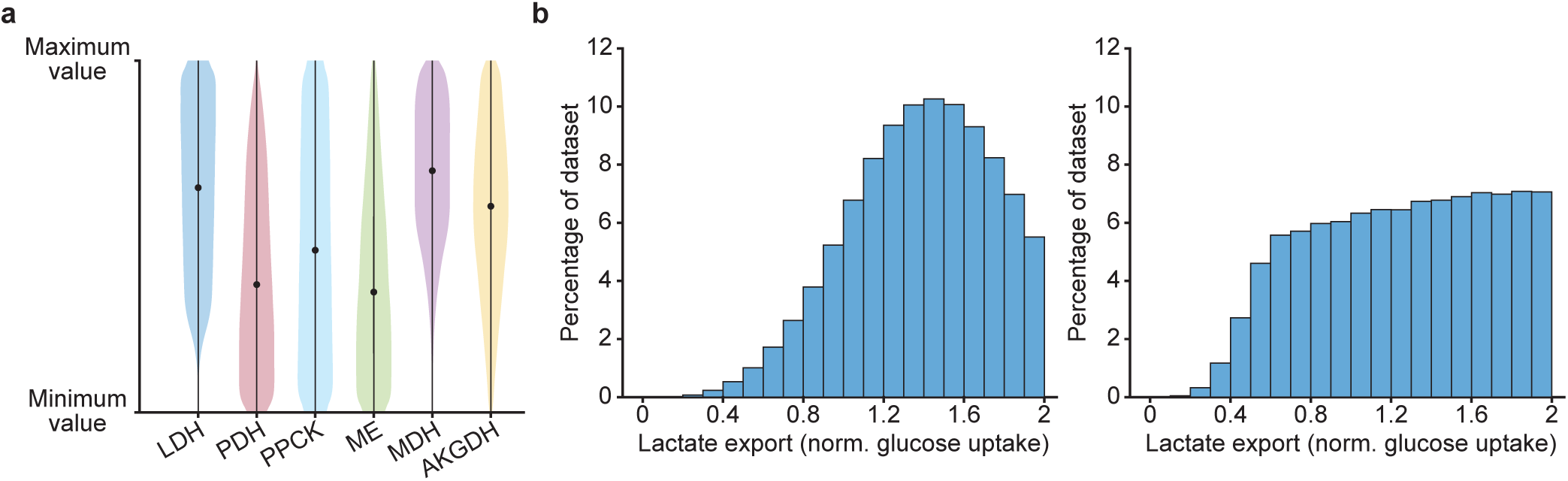
Curated sampling algorithms generate more uniform distributions of key fluxes. **a,** Key metabolic fluxes determined to have a large impact ANN model accuracy were chosen for rejection sampling after an initial artificial centering hit-and-run sampling of the constrained CCM model (see **Methods**). The resulting flux distributions were less skewed relative to the flux solution space. **b,** A representative change in the distribution of lactate export (a free flux determinant of many TCA cycle fluxes) before (left) and after (right) the rejection sampling procedure is plotted. The lack of a peak in the rejection sampled distribution represented a reduced sample bias in the training data.

## Notes

### Competing Interest Statement

The authors have declared no competing interest.

### Summary of Updates

This revision includes updated sections of text, further analyses of improved models, new statistical methods for confirming predictions, modifications to authorship, and updated figures, and expanded supplemental files.

