## Supplemental Information for "Accurate and rapid determination of metabolic flux by deep learning of isotope patterns"

Supplementary Tables 1-15

#### Supplementary Tables

#### Supplementary Table 1. List of tracers used for each metabolic model

| | | *Tracer experiments used for ML training for each metabolic model* | | | | | |
|---|---|---|---|---|---|---|---|
| **Model** | **Simple toy model** | [1-$^{13}$C$_1$]-A | [2-$^{13}$C$_1$]-A | [1,2-$^{13}$C$_2$]-A | [3-$^{13}$C$_1$]-A | [1,3-$^{13}$C$_2$]-A | [2,3-$^{13}$C$_2$]-A |
| | **Upper glycolysis** | [1,2-$^{13}$C$_2$]-glucose | | | | | |
| | **Glycolysis** | [1,2-$^{13}$C$_2$]-glucose<br>[5-D]-glucose | | | | | |
| | **GlyPPP** | [1,2,3-$^{13}$C$_3$]-glucose | [1,2-$^{13}$C$_2$]-glucose | [1-$^{13}$C$_1$]-glucose | [2,3,4,5,6-$^{13}$C$_5$]-glucose | [2,3-$^{13}$C$_2$]-glucose | [2-$^{13}$C$_1$]-glucose |
| | | [3,4-$^{13}$C$_2$]-glucose | [3-$^{13}$C$_1$]-glucose | [4,5,6-$^{13}$C$_3$]-glucose | [4,5-$^{13}$C$_2$]-glucose | [4-$^{13}$C$_1$]-glucose | [5-$^{13}$C$_1$]-glucose |
| | | [6-$^{13}$C$_1$]-glucose | [U-$^{13}$C$_6$]-glucose | [2,5,6-$^{13}$C$_3$]-glucose | [1,5,6-$^{13}$C$_3$]-glucose | [1,2,4-$^{13}$C$_3$]-glucose | [2,4-$^{13}$C$_2$]-glucose |
| | | [1,4-$^{13}$C$_2$]-glucose | [4,6-$^{13}$C$_2$]-glucose | [2,5-$^{13}$C$_2$]-glucose | [1,3-$^{13}$C$_2$]-glucose | [5,6-$^{13}$C$_2$]-glucose | [1,6-$^{13}$C$_2$]-glucose |
| | **CCM** | [U-$^{13}$C$_6$]-glucose<br>[$^{12}$C]-glutamine | [$^{12}$C]-glucose<br>[U-$^{13}$C$_5$]-glutamine | [1-$^{13}$C$_1$]-glucose<br>[$^{12}$C]-glutamine | [1-$^{13}$C$_1$]-glucose<br>[U-$^{13}$C$_5$]-glutamine | [2-$^{13}$C$_1$]-glucose<br>[$^{12}$C]-glutamine | |
| | | [2-$^{13}$C$_1$]-glucose<br>[U-$^{13}$C$_5$]-glutamine | [1,2-$^{13}$C$_2$]-glucose<br>[$^{12}$C]-glutamine | [1,2-$^{13}$C$_2$]-glucose<br>[U-$^{13}$C$_5$]-glutamine | [6-$^{13}$C$_1$]-glucose<br>[$^{12}$C]-glutamine | [6-$^{13}$C$_1$]-glucose<br>[U-$^{13}$C$_5$]-glutamine | |
| | | [1,6-$^{13}$C$_2$]-glucose<br>[$^{12}$C]-glutamine | [1,6-$^{13}$C$_2$]-glucose<br>[U-$^{13}$C$_5$]-glutamine | [3-$^{13}$C$_1$]-glucose<br>[$^{12}$C]-glutamine | [3-$^{13}$C$_1$]-glucose<br>[U-$^{13}$C$_5$]-glutamine | [5-$^{13}$C$_1$]-glucose<br>[$^{12}$C]-glutamine | |
| | | [5-$^{13}$C$_1$]-glucose<br>[U-$^{13}$C$_5$]-glutamine | [4-$^{13}$C$_1$]-glucose<br>[$^{12}$C]-glutamine | [4-$^{13}$C$_1$]-glucose<br>[U-$^{13}$C$_5$]-glutamine | [5,6-$^{13}$C$_2$]-glucose<br>[$^{12}$C]-glutamine | [5,6-$^{13}$C$_2$]-glucose<br>[U-$^{13}$C$_5$]-glutamine | |

#### Supplementary Table 2. Reactions and constraints in simple toy model

| **Reactions and constraints for simple toy metabolic model** | | | | |
|---|---|---|---|---|
| **Reaction** | **Reactants** | **Products** | **Net flux constraint** | **Exchange flux constraint** |
| v1 | A | B | = 1 | = 0 |
| v2 | B | D | | ≤ 100 |
| v3 | C | C<br>D | | = 0 |
| v4 | B<br>C | D<br>2 E | | = 0 |
| v5 | D | F | | = 0 |
| Efflux | E | | | |
| | F | | | |
| Influx | | A | = 1 | |

#### Supplementary Table 3. Reactions and constraints in upper and full glycolysis

| Reactions and constraints for upper and full glycolysis metabolic model | | | | | | | |
|---|---|---|---|---|---|---|---|
| **EC Number** | **Reaction** | **Abbreviation** | **Reactants** | **Products** | **Net flux constraint** | **Exchange flux constraint** | **Included in upper glycolysis** |
| 2.7.1.1 | hexokinase | hex | GLC | G6P | = 1 | | Y |
| 5.3.1.9 | phosphoglucose isomerase | pgi | G6P | F6P | ≥ 0<br>≤ 1.1 | ≤ 50 | Y |
| 2.7.1.11 | phosphofructokinase | pfk | F6P | FBP | | ≤ 50 | Y |
| 4.1.2.13 | fructose-bisphosphate aldolase | fba | FBP | DHAP<br>GAP | | ≤ 50 | Y |
| 5.3.1.1 | triose-phosphate isomerase | tpi | DHAP | GAP | | ≤ 50 | Y |
| 1.2.1.12 | glyceraldehyde-3-phosphate dehydrogenase | gapd | GAP | BPG | | ≤ 100 | Y |
| 2.7.2.3 | phosphoglycerate kinase | pgk | 13BPG | 3PG | | ≤ 100 | |
| 5.4.2.11 | phosphoglycerate mutase | pgm1 | 3PG | 23BPG | | ≤ 100 | |
| 3.1.3.80 | 2,3-bisphosphoglycerate 3-phosphatase | pgm2 | 23BPG | 2PG | | ≤ 100 | |
| 4.2.1.11 | enolase | eno | 2PG | PEP | | ≤ 100 | |
| 2.7.1.40 | pyruvate kinase | pyk | PEP | PYR | | = 0 | |
| | Efflux | | H | | ≥ 990<br>≤ 1000 | | Y |
| | | | NADH | | ≥ 990<br>≤ 1000 | | |
| | | | G6P | | ≥ 0.01<br>≤ 0.05 | | Y |
| | | | F6P | | ≥ 0.01<br>≤ 0.05 | | Y |
| | | | DHAP | | ≥ 0.01<br>≤ 0.05 | | Y |
| | | | 3PG | | ≥ 0.01<br>≤ 0.05 | | |
| | | | PEP | | ≥ 0.01<br>≤ 0.05 | | |
| | Influx | | | GLC | = 1 | | Y |
| | | | | H | ≥ 990<br>≤ 1000 | | Y |
| | | | | NADH | ≥ 90<br>≤ 100 | | |

### Supplementary Table 4. Reactions and constraints in glycolysis and pentose phosphate pathway

| Reactions and constraints for glycolysis and pentose phosphate pathway metabolic model | | | | | | | | | | |
|---|---|---|---|---|---|---|---|---|---|---|
| | | | | | Constraints 1 (20%) | | Constraints 2 (30%) | | Constraints 3 (50%) | |
| EC Number | Reaction | Abbreviation | Reactants | Products | Net flux constraint | Exchange flux constraint | Net flux constraint | Exchange flux constraint | Net flux constraint | Exchange flux constraint |
| 2.7.1.1 | hexokinase | hex | GLC | G6P | = 1 | | = 1 | | = 1 | |
| 5.3.1.9 | phosphoglucose isomerase | pgi | G6P | F6P | ≥ 0<br>≤ 1.1 | ≤ 0.2 | ≥ 0<br>≤ 1.1 | ≤ 2 | ≥ 0<br>≤ 1.1 | ≤ 5 |
| 2.7.1.11 | phosphofructokinase | pfk | F6P | FBP | | = 0 | | = 0 | | = 0 |
| 4.1.2.13 | fructose-bisphosphate aldolase | fba | FBP | DHAP<br>GAP | | ≤ 0.2 | | ≤ 2 | | ≤ 3 |
| 5.3.1.1 | triose-phosphate isomerase | tpi | DHAP | GAP | | ≤ 0.2 | | ≤ 2 | | ≤ 5 |
| 1.2.1.12 | glyceraldehyde-3-phosphate dehydrogenase | gapd | GAP | BPG | | ≤ 0.4 | | ≤ 4 | | ≤ 10 |
| 2.7.2.3 | phosphoglycerate kinase | pgk | 13BPG | 3PG | | ≤ 0.4 | | ≤ 4 | | ≤ 10 |
| 5.4.2.11 | phosphoglycerate mutase | Pgm1 | 3PG | 23BPG | | ≤ 0.4 | | ≤ 4 | | ≤ 10 |
| 3.1.3.80 | 2,3-bisphosphoglycerate 3-phosphatase | Pgm2 | 23BPG | 2PG | | ≤ 0.4 | | ≤ 4 | | ≤ 10 |
| 4.2.1.11 | enolase | eno | 2PG | PEP | | ≤ 0.4 | | ≤ 4 | | ≤ 10 |
| 1.1.1.49 | glucose-6-phosphate dehydrogenase | G6pdh | G6P | 6PG | ≤ 0.05 | = 0 | ≤ 0.2 | = 0 | ≤ 1 | = 0 |
| 1.1.1.44 | phosphogluconate dehydrogenase | gnd | 6PG | RU5P<br>CO2 | | ≤ 0.05 | | ≤ 0.2 | | ≤ 1 |
| 5.3.1.6 | ribose-5-phosphate isomerase | rpi | Ru5P | R5P | | ≤ 0.05 | | ≤ 0.2 | | ≤ 1 |
| 5.1.3.1 | ribulose-5-phosphate 3-epimerase | Rpe | Ru5P | Xu5P | | ≤ 0.05 | | ≤ 0.2 | | ≤ 1 |
| 2.2.1.1 | transketolase | Tkt2 | X5P<br>E4P | GAP<br>F6P | | ≤ 0.05 | | ≤ 0.2 | | ≤ 1 |
| 2.2.1.1 | transketolase | Tkt1 | X5P<br>R5P | S7P<br>GAP | | ≤ 0.05 | | ≤ 0.2 | | ≤ 1 |
| 2.2.1.2 | transaldolase | tal | GAP<br>S7P | E4P<br>F6P | | ≤ 0.05 | | ≤ 0.2 | | ≤ 1 |
| | sedoheptulose bisphosphate aldolase | sba | DHAP<br>E4P | SBP | | ≤ 0.05 | | ≤ 0.2 | | ≤ 1 |
| 3.1.3.37 | sedoheptulose-1,7-bisphosphatase | sbpase | SBP | S7P | ≥ 0.1<br>≤ 0 | ≤ 0.01 | ≥ 0.1<br>≤ 0 | ≤ 0.01 | ≥ 0.1<br>≤ 0 | ≤ 0.01 |
| | Efflux | | CO2 | | ≥ 490<br>≤ 500 | | ≥ 490<br>≤ 500 | | ≥ 490<br>≤ 500 | |
| | | | H | | ≥ 900<br>≤ 1000 | | ≥ 900<br>≤ 1000 | | ≥ 900<br>≤ 1000 | |
| | | | NADPH | | ≥ 90<br>≤ 100 | | ≥ 90<br>≤ 100 | | ≥ 90<br>≤ 100 | |
| | | | NADH | | ≥ 90<br>≤ 100 | | ≥ 90<br>≤ 100 | | ≥ 90<br>≤ 100 | |
| | | | G6P | | ≥ 0.01<br>≤ 0.05 | | ≥ 0.01<br>≤ 0.05 | | ≥ 0.01<br>≤ 0.05 | |
| | | | F6P | | ≥ 0.01<br>≤ 0.05 | | ≥ 0.01<br>≤ 0.05 | | ≥ 0.01<br>≤ 0.05 | |
| | | | DHAP | | ≥ 0.01<br>≤ 0.05 | | ≥ 0.01<br>≤ 0.05 | | ≥ 0.01<br>≤ 0.05 | |
| | | | R5P | | ≥ 0.01<br>≤ 0.05 | | ≥ 0.01<br>≤ 0.05 | | ≥ 0.01<br>≤ 0.05 | |
| | | | E4P | | ≥ 0.01<br>≤ 0.05 | | ≥ 0.01<br>≤ 0.05 | | ≥ 0.01<br>≤ 0.05 | |
| | | | 3PG | | ≥ 0.01<br>≤ 0.05 | | ≥ 0.01<br>≤ 0.05 | | ≥ 0.01<br>≤ 0.05 | |
| | | | PEP | | ≥ 0.01 | | ≥ 0.01 | | ≥ 0.01 | |
| | Influx | | | CO2 | ≥ 490<br>≤ 500 | | ≥ 490<br>≤ 500 | | ≥ 490<br>≤ 500 | |
| | | | | H | ≥ 900<br>≤ 1000 | | ≥ 900<br>≤ 1000 | | ≥ 900<br>≤ 1000 | |
| | | | | NADPH | ≥ 90<br>≤ 100 | | ≥ 90<br>≤ 100 | | ≥ 90<br>≤ 100 | |
| | | | | NADH | ≥ 90<br>≤ 100 | | ≥ 90<br>≤ 100 | | ≥ 90<br>≤ 100 | |
| | | | | GLC | = 1 | | = 1 | | = 1 | |

### Supplementary Table 5. Reactions and constraints in central carbon metabolism

| Reactions and constraints for central carbon metabolism model | | | | | | |
|---|---|---|---|---|---|---|
| EC Number | Reaction | Abbreviation | Reactants | Products | Net flux | Exchange flux |
| 2.7.1.1 | hexokinase | hex | GLC | G6P | | = 0 |
| 5.3.1.9 | phosphoglucose isomerase | pgi | G6P | F6P | ≥ 0.8<br>≤ 1 | ≤ 20 |
| 2.7.1.11 | phosphofructokinase | pfk | F6P | FBP | ≥ 0.8<br>≤ 1 | = 0 |
| 4.1.2.13 | fructose-bisphosphate aldolase | fba | FBP | DHAP<br>GAP | ≥ 0.8<br>≤ 1 | ≤ 4 |
| 5.3.1.1 | triose-phosphate isomerase | tpi | DHAP | GAP | ≥ 0.8<br>≤ 1 | ≤ 4 |
| 1.2.1.12 | glyceraldehyde-3-phosphate dehydrogenase | gapd | GAP | BPG | ≥ 1.6<br>≤ 2 | ≤ 20 |
| 2.7.2.3 | phosphoglycerate kinase | pgk | 13BPG | PGA | ≥ 1.6<br>≤ 2 | ≤ 20 |
| 4.2.1.11 | enolase | eno | PGA | PEP | ≥ 1.6<br>≤ 2 | ≤ 20 |
| 2.7.1.40 | pyruvake kinase | pyk | PEP | PYR | ≥ 1.6<br>≤ 2 | = 0 |
| 1.1.1.27 | lactate dehydrogenase | ldh | PYR | LAC | ≥ 0<br>≤ 2 | ≤ 10 |
| 1.1.1.49 | glucose-6-phosphate dehydrogenase | G6PDH | G6P | 6PG | ≥ 0<br>≤ 0.2 | = 0 |
| 1.1.1.44 | phosphogluconate dehydrogenase | gnd | 6PG | Ru5P<br>CO2 | ≥ 0<br>≤ 0.2 | ≤ 1 |
| 5.3.1.6 | ribose-5-phosphate isomerase | rpi | Ru5P | R5P | ≥ 0<br>≤ 0.1 | ≤ 2 |
| 5.1.3.1 | ribulose-5-phosphate 3-epimerase | Rpe | Ru5P | Xu5P | ≥ 0<br>≤ 0.1 | ≤ 2 |
| 2.2.1.1 | transketolase | Tkt2 | X5P<br>E4P | GAP<br>F6P | ≥ 0<br>≤ 0.1 | ≤ 1 |
| 2.2.1.1 | transketolase | Tkt1 | X5P<br>R5P | S7P<br>GAP | ≥ 0<br>≤ 0.1 | ≤ 1 |
| 2.2.1.2 | transaldolase | tal | GAP<br>S7P | E4P<br>F6P | ≥ 0<br>≤ 0.1 | ≤ 1 |
| | sedoheptulose bisphosphate aldolase | sba | DHAP<br>E4P | SBP | ≥ 0<br>≤ 0.01 | ≤ 0.5 |
| 3.1.3.37 | sedoheptulose-1,7-bisphosphatase | sbpase | SBP | S7P | ≥ 0 | = 0 |
| 4.1.1.32 | phosphoenolpyruvate carboxykinase | ppck | OAA | PEP<br>CO2 | ≥ 0<br>≤ 0.05 | = 0 |
| 1.1.1.40 | malic enzyme | me | MAL | PYR<br>CO2 | ≥ 0<br>≤ 0.5 | = 0 |
| 6.4.1.1 | pyruvate carboxylase | pc | PYR<br>CO2 | OAA | ≥ 0<br>≤ 0.3 | = 0 |
| 1.2.4.1 | pyruvate dehydrogenase | pdh | PYR | AcCoA<br>CO2 | ≥ 0<br>≤ 2 | = 0 |
| 2.3.3.1 | citrate synthase | cs | OAA<br>AcCoA | CitICit | ≤ 0.4 | = 0 |
| 2.3.3.8 | atp citrate lyase | ACITL | CitICit | OAA<br>AcCoA_cyt | < 0.1 | = 0 |
| 1.1.1.42 | isocitrate dehydrogenase | ICDH | CitICit | OGA<br>CO2 | ≤ 0.4 | ≤ 0.4 |
| 1.2.4.2 | alpha-ketoglutarate dehydrogenase | AKGDH | OGA | SuccCoA<br>CO2 | ≤ 0.6 | ≤ 0.1 |
| 6.2.1.5 | succinate synthase | SUCOAS | SuccCoA | Succ | ≤ 0.6 | = 0 |
| 1.3.5.1 | succinate dehydrogenase | SUCD | Succ | Fum | ≤ 0.6 | ≤ 5 |
| 4.2.1.2 | fumarase | FUM | Fum | MAL | ≤ 0.6 | ≤ 10 |
| 1.1.1.37 | malate dehydrogenase | MDH | MAL | OAA | ≥ –0.1<br>≤ 0.4 | ≤ 20 |
| 2.6.1.2 | alanine transaminase | PYR_Ala | PYR<br>Glu | Ala<br>OGA | ≥ 0<br>≤ 0.02 | ≤ 0.1 |
| 1.4.1.2 | glutamate dehydrogenase | OGA_Glu | OGA | Glu | ≥ –0.5<br>≤ 0.1 | ≤ 1 |
| 3.5.1.2 | glutaminase | OGA_Gln | Glu | Gln | ≤ 0 | = 0 |
| | Efflux | | CO2 | | ≥ 490<br>≤ 500 | |
| | | | G6P | | = 0.05 | |
| | | | DHAP | | = 0.01 | |
| | | | R5P | | = 0.004 | |
| | | | PGA | | = 0.01 | |
| | | | PYR | | = 0.02 | |
| | | | LAC | | ≤ 2 | |
| | | | Ala | | = 0.01 | |
| | | | AcCoA | | ≤ 0.5<br>≥ 0.001 | |
| | | | Glu | | = 0.05 | |
| | | | OAA | | = 0.05 | |
| | Influx | | | CO2 | ≥ 490<br>≤ 500 | |
| | | | | GLC | =1 | |
| | | | | GLN | ≥ 0<br>≤ 5 | |
| | | | | Ac | ≤ 0.5 | |
| | IN_Gln – EX_Gln | | | | ≥ 0<br>≤ 0.5 | |
| | EX_CO2 – IN_CO2 | | | | ≥ 0 | |

#### Supplementary Table 6. Standard errors of flux prediction from complete labeling data

**Standard errors of flux prediction from complete labeling data**

| Simple toy model | |
|---|---|
| **% Error** | |
| v2 | 1.3% |
| v3 | 0.4% |
| v4 | 0.4% |
| v5 | 0.3% |
| EX_E | 0.4% |
| EX_F | 0.3% |
| v2_xch | 1.8% |

| Upper glycolysis | |
|---|---|
| **% Error** | |
| G6P_EX | 46.2% |
| F6P_EX | 64.8% |
| DHAP_EX | 59.8% |
| PGI_xch | 2.6% |
| PFK_xch | 2.3% |
| FBA_xch | 1.8% |
| TPI_xch | 1.8% |

| Glycolysis | |
|---|---|
| **% Error** | |
| H_IN | 0.7% |
| NADH_IN | 3.0% |
| PGI | 1.5% |
| PFK | 2.0% |
| FBA | 2.0% |
| TPI | 2.3% |
| GAPD | 2.1% |
| PGK | 2.1% |
| PGM1 | 2.2% |
| PGM2 | 2.2% |
| ENO | 2.2% |
| PYK | 2.3% |
| H_EX | 0.7% |
| NADH_EX | 2.9% |
| G6P_EX | 51.9% |
| F6P_EX | 50.3% |
| DHAP_EX | 59.9% |
| PG3_EX | 74.7% |
| PEP_EX | 56.5% |
| PYR_EX | 2.3% |
| PGI_xch | 2.4% |
| PFK_xch | 1.8% |
| FBA_xch | 1.8% |
| TPI_xch | 2.9% |
| GAPD_xch | 16.5% |
| PGK_xch | 15.2% |
| PGM1_xch | 13.4% |
| PGM2_xch | 13.2% |
| ENO_xch | 10.7% |

| Glycolysis and PPP | |
|---|---|
| **Error** | |
| CO2_IN | 3.03 |
| H_IN | 3.23 |
| NADPH_IN | 3.15 |
| NADH_IN | 2.78 |
| PGI | 0.02 |
| PFK | 0.04 |
| FBA | 0.04 |
| TPI | 0.02 |
| GAPD | 0.04 |
| PGK | 0.04 |
| PGM1 | 0.05 |
| PGM2 | 0.05 |
| ENO | 0.05 |
| G6PDH | 0.02 |
| GND | 0.02 |
| RPI | 0.01 |
| RPE | 0.01 |
| TKT2 | 0.01 |
| TKT1 | 0.01 |
| TAL | 0.03 |
| SBA | 0.03 |
| SBPASE | 0.03 |
| CO2_EX | 3.03 |
| H_EX | 3.23 |
| NADPH_EX | 3.15 |
| NADH_EX | 2.78 |
| G6P_EX | 0.01 |
| F6P_EX | 0.01 |
| DHAP_EX | 0.01 |
| R5P_EX | 0.01 |
| E4P_EX | 0.01 |
| PG3_EX | 0.01 |
| PEP_EX | 0.05 |
| PGI_xch | 0.06 |
| FBA_xch | 0.11 |
| TPI_xch | 0.05 |
| GAPD_xch | 0.69 |
| PGK_xch | 0.69 |
| PGM1_xch | 0.69 |
| PGM2_xch | 0.70 |
| ENO_xch | 0.70 |
| GND_xch | 0.06 |
| RPI_xch | 0.06 |
| RPE_xch | 0.06 |
| TKT2_xch | 0.05 |
| TKT1_xch | 0.06 |
| TAL_xch | 0.06 |
| SBA_xch | 0.06 |
| SBPASE_xch | 0.00 |
| PGI_leak_xch | 0.00 |
| RPI_leak_xch | 0.00 |

| Central carbon metabolism | | | |
|---|---|---|---|
| **Error** | | | |
| IN_CO2 | 3.61 | mdh | 0.08 |
| EX_CO2 | 3.62 | PYR_Ala | 0.01 |
| hk | 0.00 | EX_PYR_Ala | 0.01 |
| pgi | 0.03 | OGA_Glu | 0.06 |
| pfk | 0.01 | IN_Gln | 1.70 |
| fba | 0.01 | IN_Glu | 0.32 |
| tpi | 0.01 | OGA_Gln | 0.32 |
| gapd | 0.01 | EX_Gln | 1.70 |
| pgk | 0.01 | IN_OAA | 0.01 |
| eno | 0.01 | EX_OGA_Glu | 0.68 |
| pyk | 0.02 | EX_AC_cyt | 0.03 |
| ldh | 0.07 | pgi_xch | 0.61 |
| EX_DHAP | 0.01 | fba_xch | 0.32 |
| EX_PGA | 0.00 | tpi_xch | 0.28 |
| EX_PYR | 0.01 | gapd_xch | 3.24 |
| EX_LAC | 0.07 | pgk_xch | 3.28 |
| EX_G6P | 0.01 | eno_xch | 2.85 |
| ppck | 0.02 | ldh_xch | 3.15 |
| me | 0.07 | ppck_xch | 0.00 |
| pc | 0.06 | me_xch | 0.00 |
| g6pdh | 0.03 | pc_xch | 0.00 |
| gnd | 0.03 | gnd_xch | 0.32 |
| rpi | 0.01 | rpi_xch | 0.63 |
| rpe | 0.02 | rpe_xch | 0.57 |
| tkt2 | 0.01 | tkt2_xch | 0.31 |
| tkt1 | 0.01 | tkt1_xch | 0.31 |
| tal | 0.01 | tal_xch | 0.19 |
| SBA | 0.00 | SBA_xch | 0.16 |
| SBPase | 0.00 | pdh_xch | 0.00 |
| EX_R5P | 0.00 | cs_xch | 0.00 |
| EX_OAA | 0.07 | icdh_xch | 0.12 |
| pdh | 0.07 | akgdh_xch | 0.03 |
| IN_AC | 0.08 | sucoas_xch | 0.00 |
| cs | 0.05 | sucd_xch | 0.45 |
| acitl | 0.03 | fum_xch | 0.73 |
| icdh | 0.05 | mdh_xch | 1.53 |
| akgdh | 0.06 | PYR_Ala_xch | 0.03 |
| sucoas | 0.06 | OGA_Glu_xch | 0.24 |
| sucd | 0.06 | OGA_Gln_xch | 0.00 |
| fum | 0.06 | | |

#### Supplementary Table 7. Standard errors of flux prediction from incomplete labeling data

**Standard errors of flux prediction from incomplete labeling data**

**Simple toy model**

| | % Error |
|---|---|
| v2 | 16.7% |
| v3 | 4.1% |
| v4 | 4.1% |
| v5 | 6.4% |
| EX_E | 4.1% |
| EX_F | 6.4% |
| v2_xch | 23.4% |

**Upper glycolysis**

| | % Error |
|---|---|
| G6P_EX | 45.8% |
| F6P_EX | 70.0% |
| DHAP_EX | 75.8% |
| PGI_xch | 11.6% |
| PFK_xch | 9.3% |
| FBA_xch | 5.2% |
| TPI_xch | 3.3% |

**Glycolysis**

| | % Error |
|---|---|
| H_IN | 0.8% |
| NADH_IN | 3.2% |
| PGI | 1.5% |
| PFK | 2.0% |
| FBA | 2.0% |
| TPI | 2.0% |
| GAPD | 1.9% |
| PGK | 1.9% |
| PGM1 | 2.0% |
| PGM2 | 2.0% |
| ENO | 2.0% |
| PYK | 2.2% |
| H_EX | 0.8% |
| NADH_EX | 3.2% |
| G6P_EX | 50.0% |
| F6P_EX | 49.2% |
| DHAP_EX | 66.9% |
| PG3_EX | 67.5% |
| PEP_EX | 60.3% |
| PYR_EX | 2.2% |
| PGI_xch | 33.0% |
| PFK_xch | 20.4% |
| FBA_xch | 11.3% |
| TPI_xch | 8.0% |
| GAPD_xch | 49.1% |
| PGK_xch | 94.6% |
| PGM1_xch | 65.6% |
| PGM2_xch | 71.7% |
| ENO_xch | 46.8% |

**Glycolysis and PPP**

| | Error |
|---|---|
| CO2_IN | 3.17 |
| H_IN | 3.35 |
| NADPH_IN | 2.88 |
| NADH_IN | 2.88 |
| PGI | 0.03 |
| PFK | 0.05 |
| FBA | 0.05 |
| TPI | 0.03 |
| GAPD | 0.05 |
| PGK | 0.05 |
| PGM1 | 0.05 |
| PGM2 | 0.05 |
| ENO | 0.05 |
| G6PDH | 0.03 |
| GND | 0.03 |
| RPI | 0.02 |
| RPE | 0.02 |
| TKT2 | 0.01 |
| TKT1 | 0.01 |
| TAL | 0.03 |
| SBA | 0.04 |
| SBPASE | 0.04 |
| CO2_EX | 3.16 |
| H_EX | 3.34 |
| NADPH_EX | 2.86 |
| NADH_EX | 2.92 |
| G6P_EX | 0.01 |
| F6P_EX | 0.01 |
| DHAP_EX | 0.01 |
| R5P_EX | 0.01 |
| E4P_EX | 0.01 |
| PG3_EX | 0.01 |
| PEP_EX | 0.05 |
| PGI_xch | 0.25 |
| FBA_xch | 0.28 |
| TPI_xch | 0.11 |
| GAPD_xch | 0.49 |
| PGK_xch | 1.06 |
| PGM1_xch | 1.12 |
| PGM2_xch | 0.85 |
| ENO_xch | 0.53 |
| GND_xch | 0.11 |
| RPI_xch | 0.04 |
| RPE_xch | 0.05 |
| TKT2_xch | 0.05 |
| TKT1_xch | 0.07 |
| TAL_xch | 0.08 |
| SBA_xch | 0.06 |
| SBPASE_xch | 0.00 |
| PGI_leak_xch | 1.15 |
| RPI_leak_xch | 2.17 |

**Central carbon metabolism**

| | Error | | Error |
|---|---|---|---|
| IN_CO2 | 3.15 | mdh | 0.10 |
| EX_CO2 | 3.14 | PYR_Ala | 0.01 |
| hk | 0.00 | EX_PYR_Ala | 0.01 |
| pgi | 0.03 | OGA_Glu | 0.07 |
| pfk | 0.01 | IN_Gln | 1.97 |
| fba | 0.01 | IN_Glu | 0.93 |
| tpi | 0.01 | OGA_Gln | 1.03 |
| gapd | 0.01 | EX_Gln | 1.62 |
| pgk | 0.01 | IN_OAA | 0.01 |
| eno | 0.01 | EX_OGA_Glu | 1.40 |
| pyk | 0.02 | EX_AC_cyt | 0.03 |
| ldh | 0.09 | pgi_xch | 5.40 |
| EX_DHAP | 0.00 | fba_xch | 1.07 |
| EX_PGA | 0.00 | tpi_xch | 0.41 |
| EX_PYR | 0.01 | gapd_xch | 6.82 |
| EX_LAC | 0.09 | pgk_xch | 6.10 |
| EX_G6P | 0.00 | eno_xch | 6.15 |
| ppck | 0.01 | ldh_xch | 3.25 |
| me | 0.09 | ppck_xch | 0.00 |
| pc | 0.07 | me_xch | 0.00 |
| g6pdh | 0.03 | pc_xch | 0.00 |
| gnd | 0.03 | gnd_xch | 0.28 |
| rpi | 0.01 | rpi_xch | 0.63 |
| rpe | 0.02 | rpe_xch | 0.55 |
| tkt2 | 0.01 | tkt2_xch | 0.31 |
| tkt1 | 0.01 | tkt1_xch | 0.31 |
| tal | 0.01 | tal_xch | 0.18 |
| SBA | 0.00 | SBA_xch | 0.16 |
| SBPase | 0.00 | pdh_xch | 0.00 |
| EX_R5P | 0.00 | cs_xch | 0.00 |
| EX_OAA | 0.08 | icdh_xch | 0.12 |
| pdh | 0.08 | akgdh_xch | 0.03 |
| IN_AC | 0.08 | sucoas_xch | 0.00 |
| cs | 0.06 | sucd_xch | 1.55 |
| acitl | 0.03 | fum_xch | 3.05 |
| icdh | 0.07 | mdh_xch | 6.30 |
| akgdh | 0.08 | PYR_Ala_xch | 0.03 |
| sucoas | 0.08 | OGA_Glu_xch | 0.31 |
| sucd | 0.08 | OGA_Gln_xch | 0.00 |
| fum | 0.08 | | |

**Supplementary Table 8. Ranking of tracers with best predictive performance for the CCM model**

| ***Ranking of tracers with best predictive performance for the CCM model*** | |
|---|---|
| Best | |
| Net intracellular fluxes | Exchange fluxes |
| [4-$^{13}C_1$]-glucose + [U-$^{13}C_5$]-glutamine | [6-$^{13}C_1$]-glucose + [U-$^{13}C_5$]-glutamine |
| [2-$^{13}C_1$]-glucose + [U-$^{13}C_5$]-glutamine | [5-$^{13}C_1$]-glucose + [$^{12}C$]-glutamine |
| [5-$^{13}C_1$]-glucoseBest[U -$^{13}C_5$]-glutamine | [6-$^{13}C_1$]-glucose + [$^{12}C$]-glutamine |
| [1,6-$^{13}C_2$]-glucose + [U-$^{13}C_5$]-glutamine | [2-$^{13}C_1$]-glucose + [$^{12}C$]-glutamine |
| [6-$^{13}C_1$]-glucose + [U-$^{13}C_5$]-glutamine | [4-$^{13}C_1$]-glucose + [$^{12}C$]-glutamine |
| [3-$^{13}C_1$]-glucose + [U-$^{13}C_5$]-glutamine | [3-$^{13}C_1$]-glucose + [U-$^{13}C_5$]-glutamine |
| [1,2-$^{13}C_2$]-glucose + [U-$^{13}C_5$]-glutamine | [1,6-$^{13}C_2$]-glucose + [$^{12}C$]-glutamine |
| [5,6-$^{13}C_2$]-glucose + [U-$^{13}C_5$]-glutamine | [1-$^{13}C_1$]-glucose + [U-$^{13}C_5$]-glutamine |
| [$^{12}C$]-glucose + [U-$^{13}C_5$]-glutamine | [1,2-$^{13}C_2$]-glucose + [$^{12}C$]-glutamine |
| [1-$^{13}C_1$]-glucose + [U-$^{13}C_5$]-glutamine | [3-$^{13}C_1$]-glucose + [$^{12}C$]-glutamine |
| [U-$^{13}C_6$]-glucose + [$^{12}C$]-glutamine | [5-$^{13}C_1$]-glucose + [U-$^{13}C_5$]-glutamine |
| [3-$^{13}C_1$]-glucose + [$^{12}C$]-glutamine | [4-$^{13}C_1$]-glucose + [U-$^{13}C_5$]-glutamine |
| [6-$^{13}C_1$]-glucose + [$^{12}C$]-glutamine | [1,2-$^{13}C_2$]-glucose + [U-$^{13}C_5$]-glutamine |
| [1-$^{13}C_1$]-glucose + [$^{12}C$]-glutamine | [1,6-$^{13}C_2$]-glucose + [U-$^{13}C_5$]-glutamine |
| [1,6-$^{13}C_2$]-glucose + [$^{12}C$]-glutamine | [2-$^{13}C_1$]-glucose + [U-$^{13}C_5$]-glutamine |
| [1,2-$^{13}C_2$]-glucose + [$^{12}C$]-glutamine | [1-$^{13}C_1$]-glucose + [$^{12}C$]-glutamine |
| [5,6-$^{13}C_2$]-glucose + [$^{12}C$]-glutamine | [5,6-$^{13}C_2$]-glucose + [$^{12}C$]-glutamine |
| [2-$^{13}C_1$]-glucose + [$^{12}C$]-glutamine | [$^{12}C$]-glucose + [U-$^{13}C_5$]-glutamine |
| [4-$^{13}C_1$]-glucose + [$^{12}C$]-glutamine | [5,6-$^{13}C_2$]-glucose + [U-$^{13}C_5$]-glutamine |
| [5-$^{13}C_1$]-glucose + [$^{12}C$]-glutamine | [U-$^{13}C_6$]-glucose + [$^{12}C$]-glutamine |
| Worst | |

**Supplementary Table 9. ANN hyperparameters**

| ANN hyperparameters | | | | | |
|---|---|---|---|---|---|
| | **Simple toy model** | **Upper glycolysis** | **Glycolysis** | **Glycolysis and PPP** | **CCM** |
| **Nodes (Input)** | 120 | 25 | 192 | 3,456 | 4,800 |
| **Nodes (Dense 1)** | 216 | 246 | 432 | 1024 | 1024 |
| **Nodes (Dense 2)** | 15 | 15 | 59 | 512 | 512 |
| **Nodes (Dense 3)** | 15 | 15 | 59 | 256 | 256 |
| **Nodes (Dense 4)** | 15 | 15 | 59 | 128 | 128 |
| **Nodes (Dense 5)** | 36 | 35 | 432 | 64 | 64 |
| **Nodes (Output)** | 6 | 7 | 18 | 33 | 55 |
| **Activation (Dense 1-5)** | RelU | | | | |
| **Loss** | MAE | MAE | MAE | Custom (see **Methods**) | Custom (see **Methods**) |
| **Optimizer** | adam | | | | |
| **Batch size** | 32 | 32 | 32 | 64 | 32 |
| **Epochs** | 1,000 | 2,000 | 2,000 | 250 | 250 |
| **Dataset size** | 100,000 | 100,000 | 100,000 | 1,000,000 | 117,077 |
| **Training:validation:testing ratio** | 80:10:10 | | | | |

**Supplementary Table 10. Proportion of data included in incomplete labeling set**

| Proportion of data in incomplete labeling set | | |
|---|---|---|
| **Model** | **Average % of data included** | **Standard deviation of % data included** |
| **Simple toy model** | 8.6% | 3.5% |
| **Upper glycolysis** | 59.8% | 22.1% |
| **Glycolysis** | 51.2% | 10.1% |
| **Glycolysis and PPP** | 2.3% | 0.4% |
| **CCM** | 2.6% | 0.4% |

### Supplementary Table 11. PCNN architecture and hyperparameters for the simple toy model

| Simple toy model PCNN architecture and hyperparameters | | | | |
|---|---|---|---|---|
| **Loss** | | MAE | | |
| **Optimizer** | | adam | | |
| **Epochs** | | 5 | | |
| **Number of masks per labeling set** | | 500 | | |
| **Dataset size** | | 100,000 | | |
| **Training:validation:testing ratio** | | 80:10:10 | | |
| **Layer name** | **Type** | **Output shape** | **Parameters** | **Connected to** |
| input_1 | InputLayer | [(None,16,16,1)] | 0 | Input |
| encoder_input | InputLayer | [(None,16,16,1)] | 0 | Input |
| conv1 | PConv2D | [(None,16,16,8),(None,16,16,8)] | 152 | input_1,encoder_input |
| batch_normalization | BatchNormalization | (None,16,16,8) | 32 | conv1 |
| tf.nn.relu | TFOpLambda | (None,16,16,8) | 0 | batch_normalization |
| conv2 | PConv2D | [(None,8,8,8),(None,8,8,8)] | 1160 | tf.nn.relu,conv1 |
| batch_normalization_1 | BatchNormalization | (None,8,8,8) | 32 | conv2 |
| tf.nn.relu_1 | TFOpLambda | (None,8,8,8) | 0 | batch_normalization_1 |
| conv3 | PConv2D | [(None,8,8,16),(None,8,8,16)] | 2320 | tf.nn.relu_1,conv2 |
| batch_normalization_2 | BatchNormalization | (None,8,8,16) | 64 | conv3 |
| tf.nn.relu_2 | TFOpLambda | (None,8,8,16) | 0 | batch_normalization_2 |
| conv4 | PConv2D | [(None,4,4,16),(None,4,4,16)] | 4624 | tf.nn.relu_2,conv3 |
| batch_normalization_3 | BatchNormalization | (None,4,4,16) | 64 | conv4 |
| tf.nn.relu_3 | TFOpLambda | (None,4,4,16) | 0 | batch_normalization_3 |
| conv5 | PConv2D | [(None,4,4,32),(None,4,4,32)] | 9248 | tf.nn.relu_3,conv4 |
| batch_normalization_4 | BatchNormalization | (None,4,4,32) | 128 | conv5 |
| tf.nn.relu_4 | TFOpLambda | (None,4,4,32) | 0 | batch_normalization_4 |
| conv6 | PConv2D | [(None,2,2,32),(None,2,2,32)] | 18464 | tf.nn.relu_4,conv5 |
| batch_normalization_5 | BatchNormalization | (None,2,2,32) | 128 | conv6 |
| tf.nn.relu_5 | TFOpLambda | (None,2,2,32) | 0 | batch_normalization_5 |
| conv7 | PConv2D | [(None,2,2,64),(None,2,2,64)] | 36928 | tf.nn.relu_5,conv6 |
| batch_normalization_6 | BatchNormalization | (None,2,2,64) | 256 | conv7 |
| tf.nn.relu_6 | TFOpLambda | (None,2,2,64) | 0 | batch_normalization_6 |
| encoder_output | PConv2D | [(None,1,1,64),(None,1,1,64)] | 73792 | tf.nn.relu_6,conv7 |
| batch_normalization_7 | BatchNormalization | (None,1,1,64) | 256 | encoder_output |
| tf.nn.relu_7 | TFOpLambda | (None,1,1,64) | 0 | batch_normalization_7 |
| up_sampling2d | UpSampling2D | (None,2,2,64) | 0 | tf.nn.relu_7 |
| up_sampling2d_1 | UpSampling2D | (None,2,2,64) | 0 | encoder_output |
| concatenate | Concatenate | (None,2,2,128) | 0 | tf.nn.relu_6,up_sampling2d |
| concatenate_1 | Concatenate | (None,2,2,128) | 0 | conv7,up_sampling2d_1 |
| conv9 | PConv2D | [(None,2,2,64),(None,2,2,64)] | 147520 | concatenate,concatenate_1 |
| batch_normalization_8 | BatchNormalization | (None,2,2,64) | 256 | conv9 |
| tf.nn.relu_8 | TFOpLambda | (None,2,2,64) | 0 | batch_normalization_8 |
| conv10 | PConv2D | [(None,2,2,32),(None,2,2,32)] | 36896 | tf.nn.relu_8,conv9 |
| batch_normalization_9 | BatchNormalization | (None,2,2,32) | 128 | conv10 |
| tf.nn.relu_9 | TFOpLambda | (None,2,2,32) | 0 | batch_normalization_9 |
| up_sampling2d_2 | UpSampling2D | (None,4,4,32) | 0 | tf.nn.relu_9 |
| up_sampling2d_3 | UpSampling2D | (None,4,4,32) | 0 | conv10 |
| concatenate_2 | Concatenate | (None,4,4,64) | 0 | tf.nn.relu_4,up_sampling2d_2 |
| concatenate_3 | Concatenate | (None,4,4,64) | 0 | conv5,up_sampling2d_3 |
| conv11 | PConv2D | [(None,4,4,32),(None,4,4,32)] | 36896 | concatenate_2,concatenate_3 |
| batch_normalization_10 | BatchNormalization | (None,4,4,32) | 128 | conv11 |
| tf.nn.relu_10 | TFOpLambda | (None,4,4,32) | 0 | batch_normalization_10 |
| conv12 | PConv2D | [(None,4,4,16),(None,4,4,16)] | 9232 | tf.nn.relu_10,conv11 |
| batch_normalization_11 | BatchNormalization | (None,4,4,16) | 64 | conv12 |
| tf.nn.relu_11 | TFOpLambda | (None,4,4,16) | 0 | batch_normalization_11 |
| up_sampling2d_4 | UpSampling2D | (None,8,8,16) | 0 | tf.nn.relu_11 |
| up_sampling2d_5 | UpSampling2D | (None,8,8,16) | 0 | conv12 |
| concatenate_4 | Concatenate | (None,8,8,32) | 0 | tf.nn.relu_2,up_sampling2d_4 |
| concatenate_5 | Concatenate | (None,8,8,32) | 0 | conv3,up_sampling2d_5 |
| conv13 | PConv2D | [(None,8,8,16),(None,8,8,16)] | 9232 | concatenate_4,concatenate_5 |
| batch_normalization_12 | BatchNormalization | (None,8,8,16) | 64 | conv13 |
| tf.nn.relu_12 | TFOpLambda | (None,8,8,16) | 0 | batch_normalization_12 |
| conv14 | PConv2D | [(None,8,8,8),(None,8,8,8)] | 2312 | tf.nn.relu_12,conv13 |
| batch_normalization_13 | BatchNormalization | (None,8,8,8) | 32 | conv14 |
| tf.nn.relu_13 | TFOpLambda | (None,8,8,8) | 0 | batch_normalization_13 |
| up_sampling2d_6 | UpSampling2D | (None,16,16,8) | 0 | tf.nn.relu_13 |
| up_sampling2d_7 | UpSampling2D | (None,16,16,8) | 0 | conv14 |
| concatenate_6 | Concatenate | (None,16,16,16) | 0 | tf.nn.relu,up_sampling2d_6 |
| concatenate_7 | Concatenate | (None,16,16,16) | 0 | conv1,up_sampling2d_7 |
| conv15 | PConv2D | [(None,16,16,8),(None,16,16,8)] | 2312 | concatenate_6,concatenate_7 |
| batch_normalization_14 | BatchNormalization | (None,16,16,8) | 32 | conv15 |
| tf.nn.relu_14 | TFOpLambda | (None,16,16,8) | 0 | batch_normalization_14 |
| decoder_output | PConv2D | [(None,16,16,1),(None,16,16,1)] | 145 | tf.nn.relu_14,conv15 |
| batch_normalization_15 | BatchNormalization | (None,16,16,1) | 4 | decoder_output |
| tf.nn.relu_15 | TFOpLambda | (None,16,16,1) | 0 | batch_normalization_15 |
| conv2d | Conv2D | (None,16,16,1) | 10 | tf.nn.relu_15 |

#### Supplementary Table 12. PCNN architecture and hyperparameters for the upper glycolysis model

| Upper glycolysis PCNN architecture and hyperparameters | | | | |
|---|---|---|---|---|
| **Loss** | | MAE | | |
| **Optimizer** | | adam | | |
| **Epochs** | | 5 | | |
| **Number of masks per labeling set** | | 500 | | |
| **Dataset size** | | 100,000 | | |
| **Training:validation:testing ratio** | | 80:10:10 | | |
| **Layer name** | **Type** | **Output shape** | **Parameters** | **Connected to** |
| input_1 | InputLayer | [(None,8,8,1)] | 0 | Input |
| encoder_input | InputLayer | [(None,8,8,1)] | 0 | Input |
| conv1 | PConv2D | [(None,8,8,4),(None,8,8,4)] | 76 | input_1,encoder_input |
| tf.nn.relu | TFOpLambda | (None,8,8,4) | 0 | conv1 |
| conv2 | PConv2D | [(None,4,4,4),(None,4,4,4)] | 292 | tf.nn.relu,conv1 |
| tf.nn.relu_1 | TFOpLambda | (None,4,4,4) | 0 | conv2 |
| conv3 | PConv2D | [(None,4,4,8),(None,4,4,8)] | 584 | tf.nn.relu_1,conv2 |
| tf.nn.relu_2 | TFOpLambda | (None,4,4,8) | 0 | conv3 |
| conv4 | PConv2D | [(None,2,2,8),(None,2,2,8)] | 1160 | tf.nn.relu_2,conv3 |
| tf.nn.relu_3 | TFOpLambda | (None,2,2,8) | 0 | conv4 |
| conv5 | PConv2D | [(None,2,2,16),(None,2,2,16)] | 2320 | tf.nn.relu_3,conv4 |
| tf.nn.relu_4 | TFOpLambda | (None,2,2,16) | 0 | conv5 |
| conv6 | PConv2D | [(None,1,1,16),(None,1,1,16)] | 4624 | tf.nn.relu_4,conv5 |
| tf.nn.relu_5 | TFOpLambda | (None,1,1,16) | 0 | conv6 |
| up_sampling2d | UpSampling2D | (None,2,2,16) | 0 | tf.nn.relu_5 |
| up_sampling2d_1 | UpSampling2D | (None,2,2,16) | 0 | conv6 |
| concatenate | Concatenate | (None,2,2,32) | 0 | tf.nn.relu_4,up_sampling2d |
| concatenate_1 | Concatenate | (None,2,2,32) | 0 | conv5,up_sampling2d_1 |
| conv7 | PConv2D | [(None,2,2,16),(None,2,2,16)] | 9232 | concatenate,concatenate_1 |
| tf.nn.relu_6 | TFOpLambda | (None,2,2,16) | 0 | conv7 |
| encoder_output | PConv2D | [(None,2,2,8),(None,2,2,8)] | 2312 | tf.nn.relu_6,conv7 |
| tf.nn.relu_7 | TFOpLambda | (None,2,2,8) | 0 | encoder_output |
| up_sampling2d_2 | UpSampling2D | (None,4,4,8) | 0 | tf.nn.relu_7 |
| up_sampling2d_3 | UpSampling2D | (None,4,4,8) | 0 | encoder_output |
| concatenate_2 | Concatenate | (None,4,4,16) | 0 | tf.nn.relu_2,up_sampling2d_2 |
| concatenate_3 | Concatenate | (None,4,4,16) | 0 | conv3,up_sampling2d_3 |
| conv9 | PConv2D | [(None,4,4,8),(None,4,4,8)] | 2312 | concatenate_2,concatenate_3 |
| tf.nn.relu_8 | TFOpLambda | (None,4,4,8) | 0 | conv9 |
| conv10 | PConv2D | [(None,4,4,4),(None,4,4,4)] | 580 | tf.nn.relu_8,conv9 |
| tf.nn.relu_9 | TFOpLambda | (None,4,4,4) | 0 | conv10 |
| up_sampling2d_4 | UpSampling2D | (None,8,8,4) | 0 | tf.nn.relu_9 |
| up_sampling2d_5 | UpSampling2D | (None,8,8,4) | 0 | conv10 |
| concatenate_4 | Concatenate | (None,8,8,8) | 0 | tf.nn.relu,up_sampling2d_4 |
| concatenate_5 | Concatenate | (None,8,8,8) | 0 | conv1,up_sampling2d_5 |
| conv11 | PConv2D | [(None,8,8,4),(None,8,8,4)] | 580 | concatenate_4,concatenate_5 |
| tf.nn.relu_10 | TFOpLambda | (None,8,8,4) | 0 | conv11 |
| decoder_output | PConv2D | [(None,8,8,1),(None,8,8,1)] | 73 | tf.nn.relu_10,conv11 |
| tf.nn.relu_11 | TFOpLambda | (None,8,8,1) | 0 | decoder_output |
| conv2d | Conv2D | (None,8,8,1) | 10 | tf.nn.relu_11 |

#### Supplementary Table 13. PCNN architecture and hyperparameters for the glycolysis model

| Glycolysis PCNN architecture and hyperparameters | | | | |
|---|---|---|---|---|
| **Loss** | MAE | | | |
| **Optimizer** | adam | | | |
| **Epochs** | 5 | | | |
| **Number of masks per labeling set** | 500 | | | |
| **Dataset size** | 100,000 | | | |
| **Training:validation:testing ratio** | 80:10:10 | | | |
| **Layer name** | **Type** | **Output shape** | **Parameters** | **Connected to** |
| input_1 | InputLayer | [(None,16,16,1)] | 0 | Input |
| encoder_input | InputLayer | [(None,16,16,1)] | 0 | Input |
| conv1 | PConv2D | [(None,16,16,4),(None,16,16,4)] | 76 | input_1,encoder_input |
| tf.nn.relu | TFOpLambda | (None,16,16,4) | 0 | conv1 |
| conv2 | PConv2D | [(None,8,8,4),(None,8,8,4)] | 292 | tf.nn.relu,conv1 |
| tf.nn.relu_1 | TFOpLambda | (None,8,8,4) | 0 | conv2 |
| conv3 | PConv2D | [(None,8,8,8),(None,8,8,8)] | 584 | tf.nn.relu_1,conv2 |
| tf.nn.relu_2 | TFOpLambda | (None,8,8,8) | 0 | conv3 |
| conv4 | PConv2D | [(None,4,4,8),(None,4,4,8)] | 1160 | tf.nn.relu_2,conv3 |
| tf.nn.relu_3 | TFOpLambda | (None,4,4,8) | 0 | conv4 |
| conv5 | PConv2D | [(None,4,4,16),(None,4,4,16)] | 2320 | tf.nn.relu_3,conv4 |
| tf.nn.relu_4 | TFOpLambda | (None,4,4,16) | 0 | conv5 |
| conv6 | PConv2D | [(None,2,2,16),(None,2,2,16)] | 4624 | tf.nn.relu_4,conv5 |
| tf.nn.relu_5 | TFOpLambda | (None,2,2,16) | 0 | conv6 |
| conv7 | PConv2D | [(None,2,2,32),(None,2,2,32)] | 9248 | tf.nn.relu_5,conv6 |
| tf.nn.relu_6 | TFOpLambda | (None,2,2,32) | 0 | conv7 |
| encoder_output | PConv2D | [(None,1,1,32),(None,1,1,32)] | 18464 | tf.nn.relu_6,conv7 |
| tf.nn.relu_7 | TFOpLambda | (None,1,1,32) | 0 | encoder_output |
| up_sampling2d | UpSampling2D | (None,2,2,32) | 0 | tf.nn.relu_7 |
| up_sampling2d_1 | UpSampling2D | (None,2,2,32) | 0 | encoder_output |
| concatenate | Concatenate | (None,2,2,64) | 0 | tf.nn.relu_6,up_sampling2d |
| concatenate_1 | Concatenate | (None,2,2,64) | 0 | conv7,up_sampling2d_1 |
| conv9 | PConv2D | [(None,2,2,32),(None,2,2,32)] | 36896 | concatenate,concatenate_1 |
| tf.nn.relu_8 | TFOpLambda | (None,2,2,32) | 0 | conv9 |
| conv10 | PConv2D | [(None,2,2,16),(None,2,2,16)] | 9232 | tf.nn.relu_8,conv9 |
| tf.nn.relu_9 | TFOpLambda | (None,2,2,16) | 0 | conv10 |
| up_sampling2d_2 | UpSampling2D | (None,4,4,16) | 0 | tf.nn.relu_9 |
| up_sampling2d_3 | UpSampling2D | (None,4,4,16) | 0 | conv10 |
| concatenate_2 | Concatenate | (None,4,4,32) | 0 | tf.nn.relu_4,up_sampling2d_2 |
| concatenate_3 | Concatenate | (None,4,4,32) | 0 | conv5,up_sampling2d_3 |
| conv11 | PConv2D | [(None,4,4,16),(None,4,4,16)] | 9232 | concatenate_2,concatenate_3 |
| tf.nn.relu_10 | TFOpLambda | (None,4,4,16) | 0 | conv11 |
| conv12 | PConv2D | [(None,4,4,8),(None,4,4,8)] | 2312 | tf.nn.relu_10,conv11 |
| tf.nn.relu_11 | TFOpLambda | (None,4,4,8) | 0 | conv12 |
| up_sampling2d_4 | UpSampling2D | (None,8,8,8) | 0 | tf.nn.relu_11 |
| up_sampling2d_5 | UpSampling2D | (None,8,8,8) | 0 | conv12 |
| concatenate_4 | Concatenate | (None,8,8,16) | 0 | tf.nn.relu_2,up_sampling2d_4 |
| concatenate_5 | Concatenate | (None,8,8,16) | 0 | conv3,up_sampling2d_5 |
| conv13 | PConv2D | [(None,8,8,8),(None,8,8,8)] | 2312 | concatenate_4,concatenate_5 |
| tf.nn.relu_12 | TFOpLambda | (None,8,8,8) | 0 | conv13 |
| conv14 | PConv2D | [(None,8,8,4),(None,8,8,4)] | 580 | tf.nn.relu_12,conv13 |
| tf.nn.relu_13 | TFOpLambda | (None,8,8,4) | 0 | conv14 |
| up_sampling2d_6 | UpSampling2D | (None,16,16,4) | 0 | tf.nn.relu_13 |
| up_sampling2d_7 | UpSampling2D | (None,16,16,4) | 0 | conv14 |
| concatenate_6 | Concatenate | (None,16,16,8) | 0 | tf.nn.relu,up_sampling2d_6 |
| concatenate_7 | Concatenate | (None,16,16,8) | 0 | conv1,up_sampling2d_7 |
| conv15 | PConv2D | [(None,16,16,4),(None,16,16,4)] | 580 | concatenate_6,concatenate_7 |
| tf.nn.relu_14 | TFOpLambda | (None,16,16,4) | 0 | conv15 |
| decoder_output | PConv2D | [(None,16,16,1),(None,16,16,1)] | 73 | tf.nn.relu_14,conv15 |
| tf.nn.relu_15 | TFOpLambda | (None,16,16,1) | 0 | decoder_output |
| conv2d | Conv2D | (None,16,16,1) | 10 | tf.nn.relu_15 |

#### Supplementary Table 14. PCNN architecture and hyperparameters for the glycolysis and pentose phosphate pathway model

| Glycolysis and pentose phosphate pathway PCNN architecture and hyperparameters | | | | |
|---|---|---|---|---|
| **Loss** | | MSE | | |
| **Optimizer** | | adam | | |
| **Epochs** | | 5 | | |
| **Number of masks per labeling set** | | 4,000 | | |
| **Dataset size** | | 1,000,000 | | |
| **Training:validation:testing ratio** | | 80:10:10 | | |
| **Layer name** | **Type** | **Output shape** | **Parameters** | **Connected to** |
| input_1 | InputLayer | [(None,80,80,1)] | 0 | Input |
| encoder_input | InputLayer | [(None,80,80,1)] | 0 | Input |
| conv1 | PConv2D | [(None,80,80,16),(None,80,80,16)] | 304 | input_1,encoder_input |
| batch_normalization | BatchNormalization | (None,80,80,16) | 64 | conv1 |
| tf.nn.relu | TFOpLambda | (None,80,80,16) | 0 | batch_normalization |
| conv2 | PConv2D | [(None,40,40,16),(None,40,40,16)] | 4624 | tf.nn.relu,conv1 |
| batch_normalization_1 | BatchNormalization | (None,40,40,16) | 64 | conv2 |
| tf.nn.relu_1 | TFOpLambda | (None,40,40,16) | 0 | batch_normalization_1 |
| conv3 | PConv2D | [(None,40,40,32),(None,40,40,32)] | 9248 | tf.nn.relu_1,conv2 |
| batch_normalization_2 | BatchNormalization | (None,40,40,32) | 128 | conv3 |
| tf.nn.relu_2 | TFOpLambda | (None,40,40,32) | 0 | batch_normalization_2 |
| conv4 | PConv2D | [(None,20,20,32),(None,20,20,32)] | 18464 | tf.nn.relu_2,conv3 |
| batch_normalization_3 | BatchNormalization | (None,20,20,32) | 128 | conv4 |
| tf.nn.relu_3 | TFOpLambda | (None,20,20,32) | 0 | batch_normalization_3 |
| conv5 | PConv2D | [(None,20,20,64),(None,20,20,64)] | 36928 | tf.nn.relu_3,conv4 |
| batch_normalization_4 | BatchNormalization | (None,20,20,64) | 256 | conv5 |
| tf.nn.relu_4 | TFOpLambda | (None,20,20,64) | 0 | batch_normalization_4 |
| conv6 | PConv2D | [(None,10,10,64),(None,10,10,64)] | 73792 | tf.nn.relu_4,conv5 |
| batch_normalization_5 | BatchNormalization | (None,10,10,64) | 256 | conv6 |
| tf.nn.relu_5 | TFOpLambda | (None,10,10,64) | 0 | batch_normalization_5 |
| conv7 | PConv2D | [(None,10,10,128),(None,10,10,128)] | 147584 | tf.nn.relu_5,conv6 |
| batch_normalization_6 | BatchNormalization | (None,10,10,128) | 512 | conv7 |
| tf.nn.relu_6 | TFOpLambda | (None,10,10,128) | 0 | batch_normalization_6 |
| encoder_output | PConv2D | [(None,5,5,128),(None,5,5,128)] | 295040 | tf.nn.relu_6,conv7 |
| batch_normalization_7 | BatchNormalization | (None,5,5,128) | 512 | encoder_output |
| tf.nn.relu_7 | TFOpLambda | (None,5,5,128) | 0 | batch_normalization_7 |
| up_sampling2d | UpSampling2D | (None,10,10,128) | 0 | tf.nn.relu_7 |
| up_sampling2d_1 | UpSampling2D | (None,10,10,128) | 0 | encoder_output |
| concatenate | Concatenate | (None,10,10,256) | 0 | tf.nn.relu_6,up_sampling2d |
| concatenate_1 | Concatenate | (None,10,10,256) | 0 | conv7,up_sampling2d_1 |
| conv9 | PConv2D | [(None,10,10,128),(None,10,10,128)] | 589952 | concatenate,concatenate_1 |
| batch_normalization_8 | BatchNormalization | (None,10,10,128) | 512 | conv9 |
| tf.nn.relu_8 | TFOpLambda | (None,10,10,128) | 0 | batch_normalization_8 |
| conv10 | PConv2D | [(None,10,10,64),(None,10,10,64)] | 147520 | tf.nn.relu_8,conv9 |
| batch_normalization_9 | BatchNormalization | (None,10,10,64) | 256 | conv10 |
| tf.nn.relu_9 | TFOpLambda | (None,10,10,64) | 0 | batch_normalization_9 |
| up_sampling2d_2 | UpSampling2D | (None,20,20,64) | 0 | tf.nn.relu_9 |
| up_sampling2d_3 | UpSampling2D | (None,20,20,64) | 0 | conv10 |
| concatenate_2 | Concatenate | (None,20,20,128) | 0 | tf.nn.relu_4,up_sampling2d_2 |
| concatenate_3 | Concatenate | (None,20,20,128) | 0 | conv5,up_sampling2d_3 |
| conv11 | PConv2D | [(None,20,20,64),(None,20,20,64)] | 147520 | concatenate_2,concatenate_3 |
| batch_normalization_10 | BatchNormalization | (None,20,20,64) | 256 | conv11 |
| tf.nn.relu_10 | TFOpLambda | (None,20,20,64) | 0 | batch_normalization_10 |
| conv12 | PConv2D | [(None,20,20,32),(None,20,20,32)] | 36896 | tf.nn.relu_10,conv11 |
| batch_normalization_11 | BatchNormalization | (None,20,20,32) | 128 | conv12 |
| tf.nn.relu_11 | TFOpLambda | (None,20,20,32) | 0 | batch_normalization_11 |
| up_sampling2d_4 | UpSampling2D | (None,40,40,32) | 0 | tf.nn.relu_11 |
| up_sampling2d_5 | UpSampling2D | (None,40,40,32) | 0 | conv12 |
| concatenate_4 | Concatenate | (None,40,40,64) | 0 | tf.nn.relu_2,up_sampling2d_4 |
| concatenate_5 | Concatenate | (None,40,40,64) | 0 | conv3,up_sampling2d_5 |
| conv13 | PConv2D | [(None,40,40,32),(None,40,40,32)] | 36896 | concatenate_4,concatenate_5 |
| batch_normalization_12 | BatchNormalization | (None,40,40,32) | 128 | conv13 |
| tf.nn.relu_12 | TFOpLambda | (None,40,40,32) | 0 | batch_normalization_12 |
| conv14 | PConv2D | [(None,40,40,16),(None,40,40,16)] | 9232 | tf.nn.relu_12,conv13 |
| batch_normalization_13 | BatchNormalization | (None,40,40,16) | 64 | conv14 |
| tf.nn.relu_13 | TFOpLambda | (None,40,40,16) | 0 | batch_normalization_13 |
| up_sampling2d_6 | UpSampling2D | (None,80,80,16) | 0 | tf.nn.relu_13 |
| up_sampling2d_7 | UpSampling2D | (None,80,80,16) | 0 | conv14 |
| concatenate_6 | Concatenate | (None,80,80,32) | 0 | tf.nn.relu,up_sampling2d_6 |
| concatenate_7 | Concatenate | (None,80,80,32) | 0 | conv1,up_sampling2d_7 |
| conv15 | PConv2D | [(None,80,80,16),(None,80,80,16)] | 9232 | concatenate_6,concatenate_7 |
| batch_normalization_14 | BatchNormalization | (None,80,80,16) | 64 | conv15 |
| tf.nn.relu_14 | TFOpLambda | (None,80,80,16) | 0 | batch_normalization_14 |
| decoder_output | PConv2D | [(None,80,80,1),(None,80,80,1)] | 289 | tf.nn.relu_14,conv15 |
| batch_normalization_15 | BatchNormalization | (None,80,80,1) | 4 | decoder_output |
| tf.nn.relu_15 | TFOpLambda | (None,80,80,1) | 0 | batch_normalization_15 |
| conv2d | Conv2D | (None,80,80,1) | 10 | tf.nn.relu_15 |

#### Supplementary Table 15. PCNN architecture and hyperparameters for the central carbon metabolism model

| Central carbon metabolism PCNN architecture and hyperparameters | | | | |
|---|---|---|---|---|
| **Loss** | MSE | | | |
| **Optimizer** | adam | | | |
| **Epochs** | 5 | | | |
| **Number of masks per labeling set** | 4,000 | | | |
| **Dataset size** | 117,077 | | | |
| **Training:validation:testing ratio** | 80:10:10 | | | |
| **Layer name** | **Type** | **Output shape** | **Parameters** | **Connected to** |
| input_1 | InputLayer | [(None,80,80,1)] | 0 | Input |
| encoder_input | InputLayer | [(None,80,80,1)] | 0 | Input |
| conv1 | PConv2D | [(None,80,80,16),(None,80,80,16)] | 304 | input_1,encoder_input |
| batch_normalization | BatchNormalization | (None,80,80,16) | 64 | conv1 |
| tf.nn.relu | TFOpLambda | (None,80,80,16) | 0 | batch_normalization |
| conv2 | PConv2D | [(None,40,40,16),(None,40,40,16)] | 4624 | tf.nn.relu,conv1 |
| batch_normalization_1 | BatchNormalization | (None,40,40,16) | 64 | conv2 |
| tf.nn.relu_1 | TFOpLambda | (None,40,40,16) | 0 | batch_normalization_1 |
| conv3 | PConv2D | [(None,40,40,32),(None,40,40,32)] | 9248 | tf.nn.relu_1,conv2 |
| batch_normalization_2 | BatchNormalization | (None,40,40,32) | 128 | conv3 |
| tf.nn.relu_2 | TFOpLambda | (None,40,40,32) | 0 | batch_normalization_2 |
| conv4 | PConv2D | [(None,20,20,32),(None,20,20,32)] | 18464 | tf.nn.relu_2,conv3 |
| batch_normalization_3 | BatchNormalization | (None,20,20,32) | 128 | conv4 |
| tf.nn.relu_3 | TFOpLambda | (None,20,20,32) | 0 | batch_normalization_3 |
| conv5 | PConv2D | [(None,20,20,64),(None,20,20,64)] | 36928 | tf.nn.relu_3,conv4 |
| batch_normalization_4 | BatchNormalization | (None,20,20,64) | 256 | conv5 |
| tf.nn.relu_4 | TFOpLambda | (None,20,20,64) | 0 | batch_normalization_4 |
| conv6 | PConv2D | [(None,10,10,64),(None,10,10,64)] | 73792 | tf.nn.relu_4,conv5 |
| batch_normalization_5 | BatchNormalization | (None,10,10,64) | 256 | conv6 |
| tf.nn.relu_5 | TFOpLambda | (None,10,10,64) | 0 | batch_normalization_5 |
| conv7 | PConv2D | [(None,10,10,128),(None,10,10,128)] | 147584 | tf.nn.relu_5,conv6 |
| batch_normalization_6 | BatchNormalization | (None,10,10,128) | 512 | conv7 |
| tf.nn.relu_6 | TFOpLambda | (None,10,10,128) | 0 | batch_normalization_6 |
| encoder_output | PConv2D | [(None,5,5,128),(None,5,5,128)] | 295040 | tf.nn.relu_6,conv7 |
| batch_normalization_7 | BatchNormalization | (None,5,5,128) | 512 | encoder_output |
| tf.nn.relu_7 | TFOpLambda | (None,5,5,128) | 0 | batch_normalization_7 |
| up_sampling2d | UpSampling2D | (None,10,10,128) | 0 | tf.nn.relu_7 |
| up_sampling2d_1 | UpSampling2D | (None,10,10,128) | 0 | encoder_output |
| concatenate | Concatenate | (None,10,10,256) | 0 | tf.nn.relu_6,up_sampling2d |
| concatenate_1 | Concatenate | (None,10,10,256) | 0 | conv7,up_sampling2d_1 |
| conv9 | PConv2D | [(None,10,10,128),(None,10,10,128)] | 589952 | concatenate,concatenate_1 |
| batch_normalization_8 | BatchNormalization | (None,10,10,128) | 512 | conv9 |
| tf.nn.relu_8 | TFOpLambda | (None,10,10,128) | 0 | batch_normalization_8 |
| conv10 | PConv2D | [(None,10,10,64),(None,10,10,64)] | 147520 | tf.nn.relu_8,conv9 |
| batch_normalization_9 | BatchNormalization | (None,10,10,64) | 256 | conv10 |
| tf.nn.relu_9 | TFOpLambda | (None,10,10,64) | 0 | batch_normalization_9 |
| up_sampling2d_2 | UpSampling2D | (None,20,20,64) | 0 | tf.nn.relu_9 |
| up_sampling2d_3 | UpSampling2D | (None,20,20,64) | 0 | conv10 |
| concatenate_2 | Concatenate | (None,20,20,128) | 0 | tf.nn.relu_4,up_sampling2d_2 |
| concatenate_3 | Concatenate | (None,20,20,128) | 0 | conv5,up_sampling2d_3 |
| conv11 | PConv2D | [(None,20,20,64),(None,20,20,64)] | 147520 | concatenate_2,concatenate_3 |
| batch_normalization_10 | BatchNormalization | (None,20,20,64) | 256 | conv11 |
| tf.nn.relu_10 | TFOpLambda | (None,20,20,64) | 0 | batch_normalization_10 |
| conv12 | PConv2D | [(None,20,20,32),(None,20,20,32)] | 36896 | tf.nn.relu_10,conv11 |
| batch_normalization_11 | BatchNormalization | (None,20,20,32) | 128 | conv12 |
| tf.nn.relu_11 | TFOpLambda | (None,20,20,32) | 0 | batch_normalization_11 |
| up_sampling2d_4 | UpSampling2D | (None,40,40,32) | 0 | tf.nn.relu_11 |
| up_sampling2d_5 | UpSampling2D | (None,40,40,32) | 0 | conv12 |
| concatenate_4 | Concatenate | (None,40,40,64) | 0 | tf.nn.relu_2,up_sampling2d_4 |
| concatenate_5 | Concatenate | (None,40,40,64) | 0 | conv3,up_sampling2d_5 |
| conv13 | PConv2D | [(None,40,40,32),(None,40,40,32)] | 36896 | concatenate_4,concatenate_5 |
| batch_normalization_12 | BatchNormalization | (None,40,40,32) | 128 | conv13 |
| tf.nn.relu_12 | TFOpLambda | (None,40,40,32) | 0 | batch_normalization_12 |
| conv14 | PConv2D | [(None,40,40,16),(None,40,40,16)] | 9232 | tf.nn.relu_12,conv13 |
| batch_normalization_13 | BatchNormalization | (None,40,40,16) | 64 | conv14 |
| tf.nn.relu_13 | TFOpLambda | (None,40,40,16) | 0 | batch_normalization_13 |
| up_sampling2d_6 | UpSampling2D | (None,80,80,16) | 0 | tf.nn.relu_13 |
| up_sampling2d_7 | UpSampling2D | (None,80,80,16) | 0 | conv14 |
| concatenate_6 | Concatenate | (None,80,80,32) | 0 | tf.nn.relu,up_sampling2d_6 |
| concatenate_7 | Concatenate | (None,80,80,32) | 0 | conv1,up_sampling2d_7 |
| conv15 | PConv2D | [(None,80,80,16),(None,80,80,16)] | 9232 | concatenate_6,concatenate_7 |
| batch_normalization_14 | BatchNormalization | (None,80,80,16) | 64 | conv15 |
| tf.nn.relu_14 | TFOpLambda | (None,80,80,16) | 0 | batch_normalization_14 |
| decoder_output | PConv2D | [(None,80,80,1),(None,80,80,1)] | 289 | tf.nn.relu_14,conv15 |
| batch_normalization_15 | BatchNormalization | (None,80,80,1) | 4 | decoder_output |
| tf.nn.relu_15 | TFOpLambda | (None,80,80,1) | 0 | batch_normalization_15 |
| conv2d | Conv2D | (None,80,80,1) | 10 | tf.nn.relu_15 |
